# On the Optimal Temporal Resolution for Information Representation in Neural Activity: A Theoretical Analysis

**DOI:** 10.64898/2026.05.19.726394

**Authors:** H. Fareed Ahmed, Toktam Samiei, Erfan Nozari

## Abstract

**Introduction:** Although neural activity is organized across multiple temporal and spatial scales, the principles determining information representation across scales remain unclear. In particular, while recent empirical results have reported mesoscale optimality in neural decoding, no theoretical accounts exist that can explain when and why such intermediate scales emerge as optimal. Here, we develop an analytical framework to determine optimal temporal scales of neural information representation and their dependence on signal and noise dynamics.

**Materials and Methods:** We formulate a multiscale model where neural population activity is represented by temporally encoded trial vectors at micro-, coarse meso-, fine meso- and macroscale resolutions. Neural responses are modeled as stimulus-dependent mean activations corrupted by temporally correlated noise, with signal and noise autocorrelation decay rates varied parametrically. Representational quality is quantified using the sensitivity index (d-prime), measuring the ability of an optimal decoder to distinguish stimulus conditions.

**Results:** We derive closed-form expressions for the sensitivity index at each temporal scale and identify signal and noise autocorrelations as key determinants of decodability. We then validate our theoretical predictions against empirical decodability estimates from synthetic neural data. Comparing these expressions under various combinations of signal and noise autocorrelations across time reveals two main regimes. First, when signal and noise correlations are absent or persistent over time, the optimal resolution falls at one of the two extremes: macroscale (resp. microscale) if signal autocorrelations are significantly stronger (resp. weaker) than noise autocorrelations. When both signal and noise autocorrelations decay, temporal integration creates a trade-off: moderate integration improves decodability by suppressing noise while preserving coherent signal, whereas excessive integration degrades signal and decodability. Therefore, only in the latter regime, mesoscale representations emerge as the optimal regime across a broad range of biologically plausible parameters.

**Discussion:** This work provides a theoretical explanation for how optimal temporal scales depend on the interplay between signal and noise autocorrelations. The framework establishes temporal integration as a principled mechanism linking multiscale neural dynamics to information representation, explains when preprocessing operations such as binning and smoothing enhance or degrade decodability, and provides testable predictions across recording modalities and neural systems.

## INTRODUCTION

Neural activity evolves across a wide range of spatial and temporal scales, from millisecond spiking in individual neurons to slowly varying dynamics measured over large populations (Buzsáki, 2006; Bressler and Menon, 2010; Breakspear, 2017). This multiscale organization is one of the defining complexities of the brain, and understanding how information is distributed, transformed, and preserved across these spatiotemporal scales remains a fundamental problem in neuroscience. A central challenge is to determine how these scales are related, how information is represented differently at each scale, and which scale most effectively captures behaviorally relevant neural information.

Perhaps the most fundamental operation linking across spatiotemporal scales is averaging (integration). At the microscale, individual neurons integrate inputs across thousands of synapses, effectively averaging presynaptic activity over space, while membrane and synaptic time constants average activity over time (Cash and Yuste, 1999; Kandel et al., 2013). At larger temporal scales, fast spiking activity gives rise to slower collective dynamics such as low-frequency oscillations, which reflect the integrated behavior of neural circuits over extended time windows (Buzsáki, 2006). At larger spatial scales, coordinated population activity emerges from the interactions of many neurons, producing meso- and macroscale signals measurable at the level of local populations, cortical areas, or whole-brain recordings (Bressler and Menon, 2010; Breakspear, 2017). In all cases, integration serves as a biological bridge across scales, transforming fine-grained neuronal events into coarser yet often more stable representations of neural activity. As a fundamental operation embedded in neural processing itself, understanding integration is therefore crucial for studying the brain and interpreting brain-related signals.

Averaging is also a central tool in neural data analysis. Most preprocessing methods—including firing-rate estimation, peri-stimulus time histograms (PSTHs), low-pass filtering, dimensionality reduction, and population summaries—rely on averaging to improve signal-to-noise ratio and/or make neural activity more interpretable (Poldrack et al., 2011; Luck, 2014; Widmann et al., 2015). At the same time, the temporal resolution at which neural activity is analyzed is itself known to influence the amount of information that can be extracted from neural responses (**??**). Recent experimental work has further suggested that neural populations may dynamically organize activity into temporally structured segments, achieving a balance between temporal integration and segregation that supports efficient sensory information representation (**?**). By suppressing trial-to-trial variability and reducing dimensionality, averaging *can* reveal structure that is otherwise obscured in noisy measurements.

Nevertheless, averaging can also blur fine temporal structure, wash out meaningful nonlinearities, and erase distinctions that are essential to the neural code. More generally, averaging involves an inevitable loss of information, as formalized by the data processing inequality (Cover and Thomas, 2006). Consistent with this perspective, information-theoretic studies have shown that the amount of stimulus information recovered from neural activity depends on the temporal precision at which spike trains are represented and decoded (**??**). Recent work has further shown that averaging can exert a strong linearizing effect, transforming functionally relevant nonlinearities into what appears as noise in macroscale measurements (Nozari et al., 2024), a finding that was later theoretically demonstrated by Ahmed et al. (2025) and Ahmed and Nozari (2023). Consequently, understanding how averaging reshapes signal and noise is not only a theoretical question about neural coding, but also a practical question about whether common preprocessing pipelines preserve or obscure behaviorally relevant information.

Specifically, the effect of temporal averaging depends primarily on the strength, distribution, and decay rate of correlations, *in both signal and noise*, over time (i.e autocorrelations). This in turn suggests that the optimal temporal scale of neural representation should be governed by the correlation structure of the underlying activity. If signal remains coherent across nearby time points while noise decorrelates rapidly, temporal integration should improve representational quality by averaging-out noise faster than it averages-out signal. In contrast, averaging can lead to a rapid loss of information if nearby bins carry dissimilar signal, thereby combining weakly related or incompatible information into a single representation. As we demonstrated in Samiei et al. (2024) using large-scale NeuroPixels recording in mice, biological population spiking activity appears to lie in the middle, with task-related information best decoded from *mesoscale* representations. As with empirical studies in general, however, the findings of this study were limited to the animal model, brain regions, task conditions, and data modality that were used therein.

In this study, we develop a general theoretical framework for identifying the optimal temporal scale of information representation in neural activity as a function of the underlying statistical structure of signal and noise across time. Using a sensitivity index that captures within-class consistency, between-class separability, and trial-to-trial variability, we derive closed-form expressions for representational discriminability across scales, and use the latter to compare the accuracy of neural representations at micro-,coarse meso-, fine meso- and macroscales. We further validate the theoretical predictions by comparing the derived sensitivity index expressions with empirical estimates obtained from synthetic neural data.

Our analysis shows that the biological consequences of averaging depend critically on how signal and noise autocorrelations decay (or not) over time. If signal remains statistically persistent across time while noise autocorrelations decay, temporal integration favors the extremes—either microscale or macroscale, depending on the decay rate of noise autocorrelations. In contrast, when both signal and noise correlations decay exponentially with temporal distance, averaging generates a nontrivial trade-off, such that limited integration strengthens coherent signal and suppresses variability, whereas excessive integration eventually leads to an auto-cancellation of both signal and noise. Under these conditions, mesoscale representations can indeed emerge as the optimal regime, as observed empirically in Samiei et al. (2024).

More broadly, this work provides a theoretical foundation for interpreting temporal averaging as a biologically meaningful operation rather than a purely technical preprocessing step. It reframes the problem of neural coding across scales in terms of how information transforms under integration, and shows that the benefits and costs of averaging can be quantified through the interplay of temporally structured signal and noise. In doing so, it offers an analytical explanation for why intermediate temporal resolutions may be privileged in neural systems, and provides a principled framework for linking multiscale neural dynamics to information representation.

## RESULTS

Neural activity can be represented at multiple temporal resolutions, ranging from fine-grained (microscale) to fully aggregated (macroscale). As noted, we hypothesize that the optimal temporal scale emerges from a trade-off between the effects of averaging on signal and noise, governed by the structure of signal and noise autocorrelations. To formalize and test this hypothesis, we develop a multiscale theoretical model that enables direct comparison of information representation across temporal resolutions, as described next.

### Theoretical framework for identifying optimal temporal resolution of neural activity

#### Context

We begin by developing a framework for quantifying the effect of averaging on neural representations in the context of population coding of an extrinsic variable. Assume we have recorded the activity of *N* neurons at a fixed microscale temporal resolution for a task with two conditions *c* = 1, 2 and fixed trial duration. Let *b* denote the total number of microscale time bins that tile the duration of each trial, and the *N* -dimensional *random vectors* 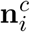, *i* = 1, 2, …, *b, c* = 1, 2 denote the population activations given each condition *c*. These can represent, e.g., the binary spiking activity of a population of neurons recorded via high density probes during 250ms trials and binned at 1ms resolution (*b* = 250), as in our prior work (Samiei et al., 2024). The two task conditions *c* = 1 and *c* = 2 can represent two distinct sensory stimuli presented to the subject, or two disjoint groups of such stimuli, and we measure the amount of information present in 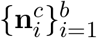 by the accuracy of an optimal decoder that seeks to decode the extrinsic task condition based on the intrinsic neural activity 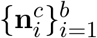.

In this context, we measure and compare the effect of averaging on neural information by comparing the decoding accuracy of four classifiers, as follows (cf. Figure 1). At the microscale, the classifier receives a realization of all population activity vectors

**Figure 1.**
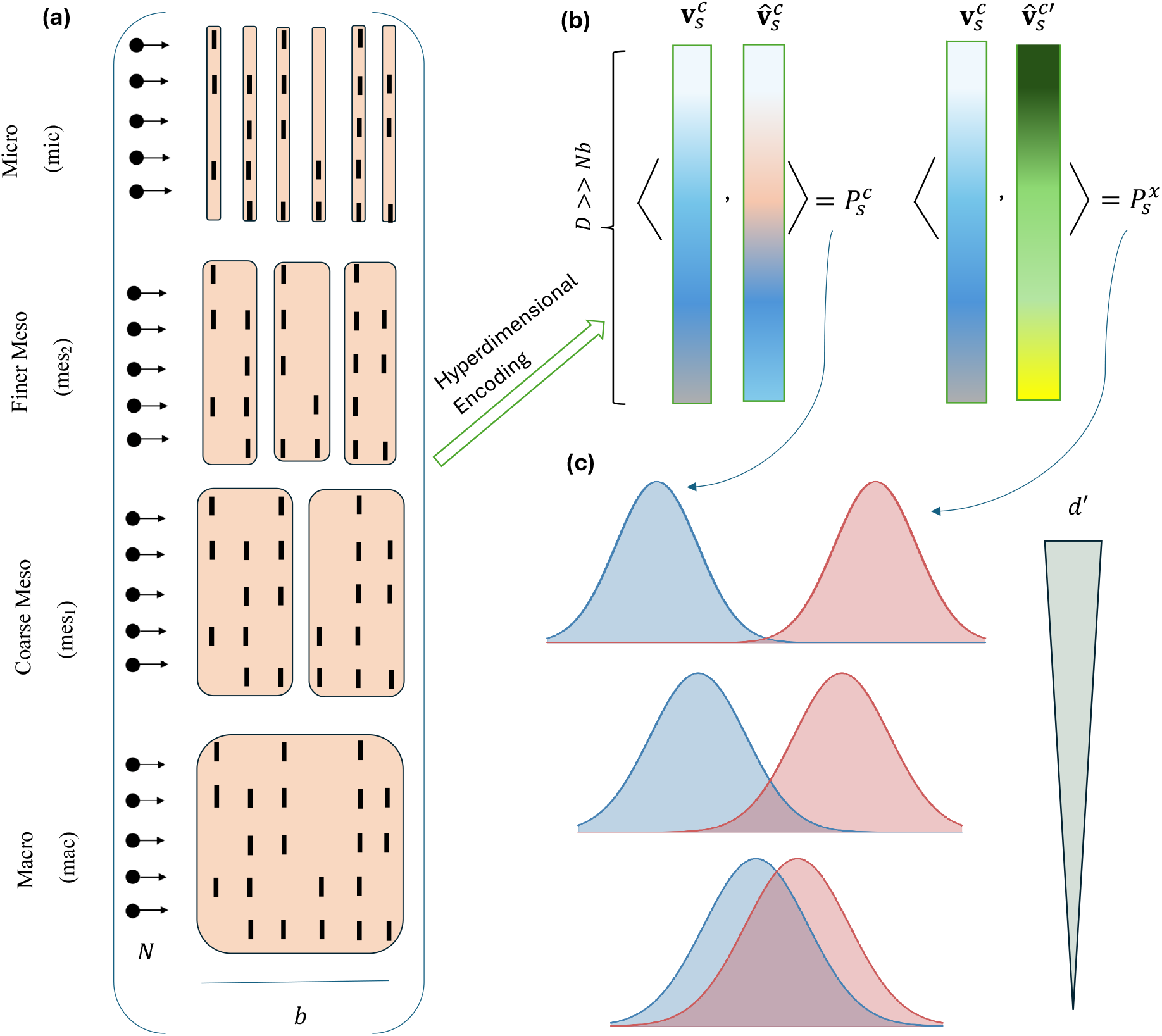
Schematic of the computational framework used for relating temporal scale and information content in neural representations. **(a)** Spiking activity of *N* hypothetical neurons is shown over *b* microscale time bins and represented (integrated) at four different temporal resolutions. Each scale of temporal integration is illustrated by colored blocks, each forming one bin at the respective scale. **(b)** The temporally integrated neural activity at each scale is hyperdimensionally encoded into a *D*-dimensional representation called a trial hypervector (Eq. (2)). The similarities of pairs of trial hypervectors are then computed using inner products, both for the case pairs that belong to the same class (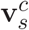 and 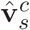) and those that belong to two different classes (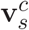 and 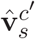), and referred to as 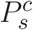 and 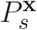, respectively. **(c)** The amount of overlap between the distributions of 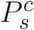 and 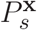, measured by the sensitivity index *d*^′^, determine the ability of a normative decoder to decode stimulus condition *c* from neural representations at that scale.

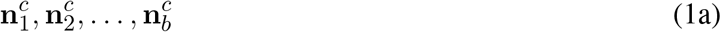

in order to predict the task condition *c*. At the opposite macroscale extreme, the classifier receives only the fully integrated activity vector

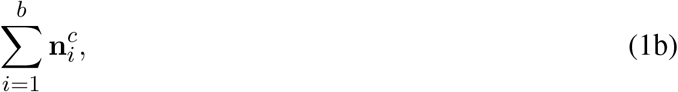

to make the same prediction. At the intermediate mesoscale we provide the decoder two alternative cases: a *coarse mesoscale* where the decoder receives the two integrated vectors

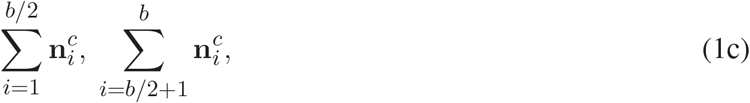

and a *fine mesoscale* in which the decoder receives *b/*2 slightly integrated vectors

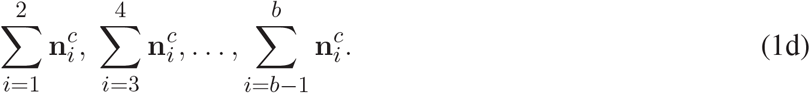

Clearly, while the two extremes are uniquely defined, Eq. (1c) and Eq. (1d) serve as representatives of the various intermediate scales. At each scale, we will quantify the accuracy of the optimal (Bayes) classifier and determine the scale that achieves maximum accuracy as a function of the signal and noise autocorrelations in the data.

#### Decoder

A unique feature, and challenge, in comparing decoding accuracies across scales is the potentially confounding effects of dimensionality. Unlike most machine learning problems where feature dimensions are either fixed or variable independently of the choice of model (due to missing data, e.g.), here the dimension of features available for decoding varies significantly and systematically with scale. As discussed in detail in Samiei et al. (2024), most modern machine learning classifiers are highly sensitive to input dimensionality, often preferring medium to small input dimensions for the same amount of information present. Therefore, to decouple differences in information content from differences in dimensionality, we first encode (lift) available information at all scales to the same hyperdimension via a lossless linear encoding. As shown by Samiei et al. (2024), this can be achieved, at the microscale, by encoding the raw neural information of Eq. (1a) into the *trial hypervector*

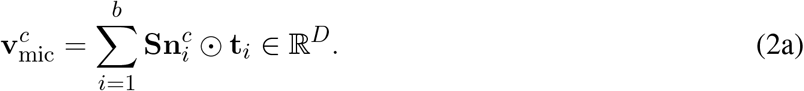

In this expression **S** ∈ ℝ^*D*×*N*^ is a matrix of spatial hypervectors that maps neural activity into a hyperdimensional space ℝ^*D*^, *D* ≫ *Nb*. In other words, each column of **S** is a *D*-dimensional spatial hypervector assigned to (and preserving the identity of the spikes of) the corresponding neuron, and the vector 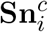 provides the superposition of all such hypervectors weighted by the activations of their corresponding neurons. Similarly, the vectors **t**_*i*_ ∈ ℝ^*D*^ denote temporal hypervectors that assign a distinct identity to each microscale bin, and the element-wise product ⊙ ‘binds’ (associates) the spatially-encoded hypervector 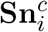 with the temporal hypervector **t**_*i*_. The final summation ‘bundles’ the resulting hypervectors over time, forming a single trial hypervector that contains information from all 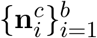. As shown in Samiei et al. (2024, Supp. Note 1), Eq. (2a) constitutes a lossless mapping such that all 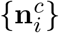 can be recovered from 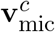 if the encoding (spatial and temporal) hypervectors are properly chosen and *D* is sufficiently large (*D* → ∞ for ideal reconstruction).

Throughout this work we assume that the encoding hypervectors are properly chosen (e.g., consisting of independent and identically distributed elements drawn from a zero-mean distribution (Samiei et al., 2024) and fixed, and can therefore be treated as deterministic. If so, as *D* → ∞, such hypervectors become asymptotically orthogonal. This implies, in particular, that **S**^⊤^**S** ≈ *γI*_*N*_, where *γ* ∈ ℝ is a constant (squared norm of each spatial hypervector).

Similarly, the raw information in Eq. (1b), Eq. (1c) and Eq. (1d) are encoded into same-dimensional trial hypervectors

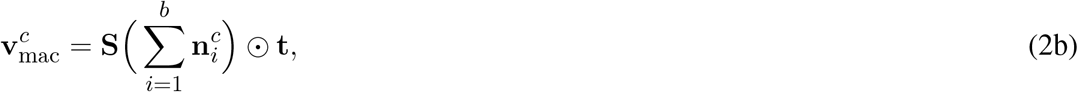

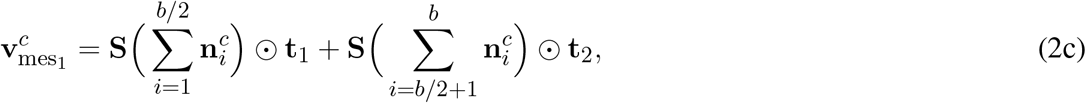

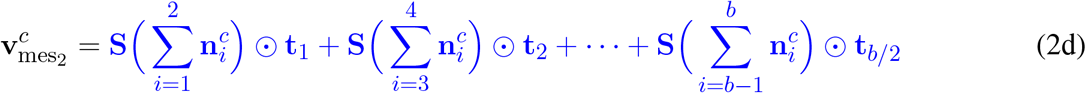

respectively. Similar to the hyperdimensional encodings at the microscale, **t** denotes the temporal hypervector associated with the full (integrated) trial at the macroscale, while **t**_1_, **t**_2_ and **t**_1_, **t**_2_, …, **t**_*b/*2_ are near-orthogonal hypervectors associated with each mesoscale bin at the two intermediate resolutions, respectively.

It is well-known that the encoding of information into hyperdimension also simplifies classification and allows for pattern recognition via simple inner products (Kanerva, 2009; Rahimi and Recht, 2007; Cortes and Vapnik, 1995). As such, let *s* ∈ *S* = {mac, mes_1_, mes_2_, mic} denote any of the four resolutions, *c* ∈ {1, 2} denote either condition, and 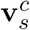 and 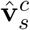 denote two independent and identically distributed (i.i.d.) trial hypervectors from the same distribution corresponding to condition *c*. Then, regardless of the exact decoding architecture used, decoding accuracy is fundamentally limited by the *within-class* and *between-class* similarities

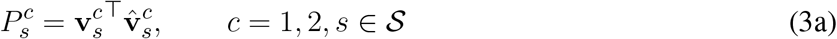

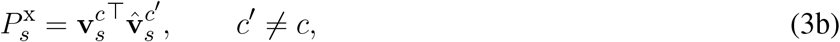

respectively (Figure 1b). Note that both 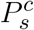 and 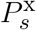 are random variables (inheriting their randomness from that of 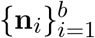), and the amount of separation between their distributions controls how accurately one can distinguish between the two conditions. If trial hypervectors are highly similar within each condition (large 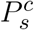) but not across conditions (small 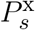), decoding accuracy will be high. If, on the other hand, trial hypervectors are equally similar to each other within and between conditions, decoding accuracy will be at chance.

Accordingly, we will measure the decodability of conditions at each scale via the sensitivity index

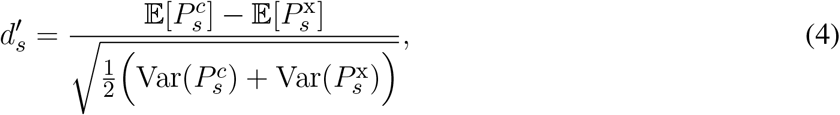

where E[·] and Var(·) denote mean and variance, respectively. We use d-prime instead of the decoding accuracy of any specific classifier given the former’s direct relation to the accuracy of an ideal (Bayes’ optimal) decoder (Averbeck and Lee, 2006). Figure 1c provides a schematic illustration of *d*^′^ and how its variation corresponds to different levels of separability between the distributions of within-class and between-class similarities. In what follows, we will derive mathematical expressions for 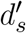 at each scale *s* ∈ *S* and study how they are related to each other depending on the correlation structures present in the data.

#### Statistical Model

As noted in the Introduction, we hypothesize that the effect of averaging on neural information depends on the relative strength of temporal autocorrelations in both the signal (condition-relevant information) and noise (condition-irrelevant variability). To formally separate the two, let

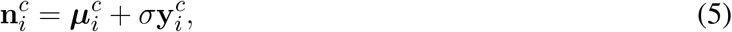

where random vector 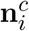 denotes the raw neural activity vector at time *i* under condition 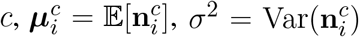 for all *i* = 1, …, *b*, and 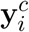 is the remaining normalized zero-mean variability in 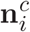 (not necessarily Gaussian). In what follows, we will refer to 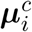 and 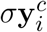 as the *signal* and *noise* components of 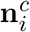, respectively.

Throughout this work we assume that noise autocorrelation decays exponentially with temporal distance. In particular, let 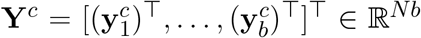 represent the concatenation of all normalized noise vectors across the trial. We assume

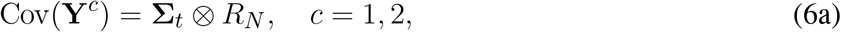

where ⊗ denotes the Kronecker product, *R*_*N*_ represents the *N* × *N* matrix of correlations among neurons at the same time, and **Σ**_*t*_ denotes the *b* × *b* matrix of exponentially decaying noise correlations within the same neuron over time, namely

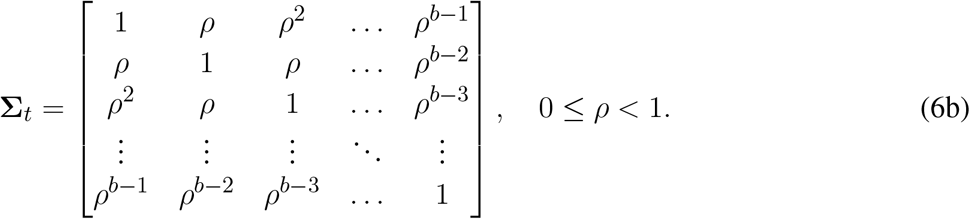

The parameter *ρ* denotes the decay rate of noise autocorrelation over time, and constitutes one of the key parameters in determining the optimal resolution for information decoding. Throughout this work we will assume the noise to be independent across the two conditions, i.e., Cov(**Y**^1^, **Y**^2^) = **0**.

Equally important in the analyses of optimal temporal resolutions is the decay rate of *signal autocorrelations* across time. We characterize this temporal autocorrelation by the cosine similarity 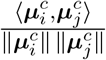 between the signal components 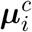. In particular, we consider two main scenarios, namely, one in which signal correlations persist throughout the trial duration

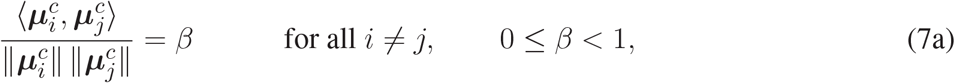

and an opposite scenario when signal auto-correlations decay with temporal distance similar to Eq. (6),

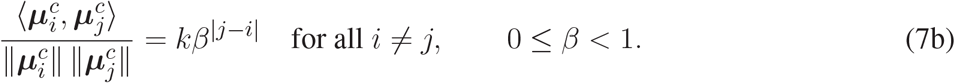

In both scenarios, the parameter *β* plays the dual role to *ρ*. Further, in Eq. (7b) *k* captures a potentially instantaneous decay in signal autocorrelation as one goes from lag 0 (*i* = *j*) to lag 1 (|*j* − *i*| = 1).

Together, this formulation enables a systematic analysis of how temporal aggregation influences discriminability across multiple scales, with the parameters *ρ* and *β* governing the temporal structure of noise and signal, respectively, and, in turn, the effect of averaging on neural information representations.

### Information Representation at the Microscale

We begin the mathematical analysis of neural information content and decodability across temporal scales by deriving a mathematical expression for sensitivity index (d-prime) in Eq. (4) at the finest scale. To enable a tractable analytical characterization of the inner-product statistics, we introduce the following assumption, which will be maintained throughout the subsequent analysis.

#### Assumption 1.

*The signal and noise components of neural responses are uncorrelated, i*.*e*.,

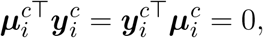

*for all time points i and conditions c*. □

Assumption 1 formulates the intuition that the task-*relevant* and task-*irrelevant* components of neural responses vary independently across the population. In the results that follow, this assumption further simplifies analytical expressions by allowing inner-product statistics to decompose into signal and noise contributions. That said, similar results can still be derived without this assumption, as shown in Supplementary Note 2 for a simple case.

We are now ready to state our first main result, providing a simple analytical expression for the sensitivity index (d-prime) at the microscale. The full proof of the result is deferred to Supplementary Note 1, with a proof sketch provided here for the interested reader.

#### Theorem 1

(Sensitivity Index at the Microscale). *Let* 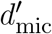 *denote the sensitivity index defined in Eq. (4) at the microscale, with similarity statistics* 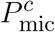 *and* 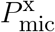 *defined in Eq. (3), trial hypervectors* 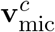 *defined in Eq. (2a), and neural responses following the statistical model in Eq. (5). Assume that Assumption 1 holds. Then*,

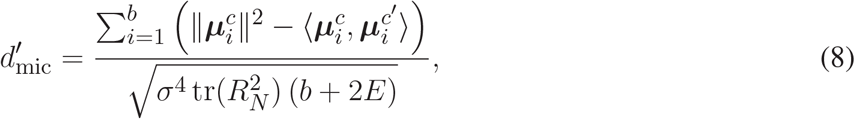

*where*

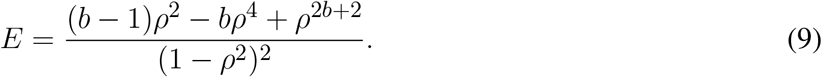

Proof sketch. In the interest of brevity we only outline the main steps of the derivation here; for the full proof see Supplementary Note 1. The core idea is that, at the microscale, the orthogonality of the temporal basis vectors **t**_*i*_ and the columns of the spatial hypervectors matrix **S**, allows the inner products between trial representations to decompose into a sum of independent contributions across microscale temporal bins. Using the decomposition in Eq. (5),

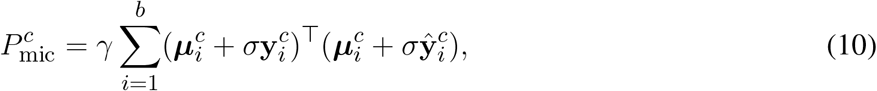

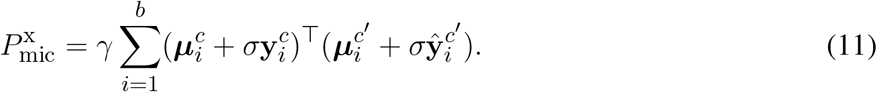

Using the statistical properties of trial hypervectors and their components, the expectations of within-class and between-class similarities reduce to,

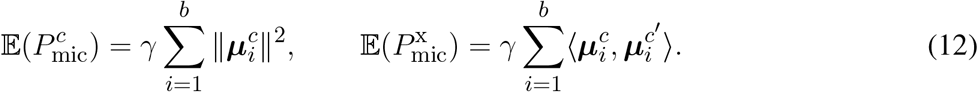

The variance terms in Eq. (4) depend, expectedly, on the noise terms in Eq. (5). Using the structure of the covariance matrix in Eq. (6), i.e., 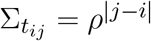, we obtain:

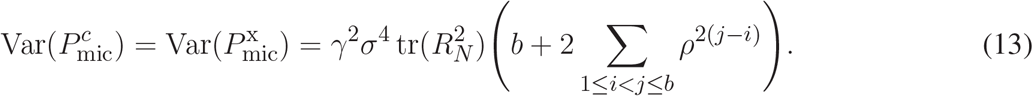

Evaluating the finite sum in Eq. (13) yields the closed-form expression for *E* in Eq. (9), and substituting these into the definition of 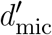 in Eq. (4) leads to the closed-form expression in Eq. (8). □

It is instructive to analyze the structure of 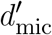 in Eq. (8) and what it reveals about information representation at the microscale. At this scale, 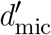 incorporates independent contributions across microscale bins, wherein the numerator of Eq. (8) represents the total integrated signal evidence accumulated through temporal summation. This reflects a setting where information is encoded locally in time, without further interactions between different temporal components. In contrast, the denominator of Eq. (8) represents integrated noise variance, incorporating temporal autocorrelations through the factor *b* + 2*E* and thus indicating that variability is not purely independent, but rather spreads across time. As a result, microscale representations exhibit noise contributions that scale linearly with the number of time points as the orthogonal basis vectors **t**_*i*_ prevent the cross-temporal coupling of trial-to-trial fluctuations, while signal remains strictly additive and localized.

Theorem 1 characterizes the decodability of information representations at the finest temporal resolution, where each microscale bin contributes independently. In the following section, we extend this analysis to coarser temporal scales, deriving analogous expressions for mesoscale and macroscale representations obtained through temporal aggregation. We will subsequently compare these results and study the conditions under which each scale provides the most amount of task-relevant information.

### Information Representation at Integrated Scales

Building on the microscale analysis described above, we next consider information representation at the *fine mesoscale*, obtained by the minimal temporal aggregation of pairs of adjacent microscale bins. The following result provides a closed-form expression for the sensitivity index 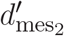, clarifying the precise contributions of signal and noise components to information decodability. The proof sketch is similar to that of Theorem 1, and the full proof is provided in Supplementary Note 1.

#### Theorem 2

(Sensitivity Index at the Fine Mesoscale). *Let* 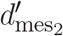 *denote the sensitivity index defined in Eq. (4), with similarity statistics* 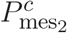 *and* 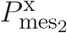 *defined in Eq. (3), trial hypervectors* 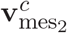 *defined in Eq. (2d), and neural responses following the statistical model in Eq. (5). Assume that Assumption 1 holds. Then*,

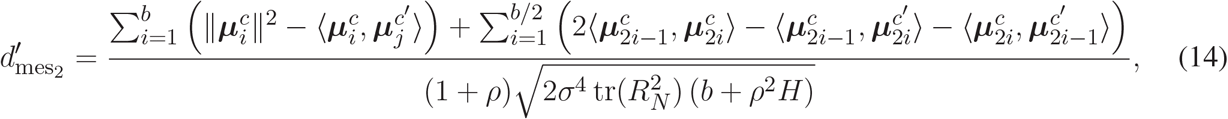

*where*

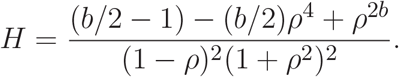

Compared to the microscale, the structure of 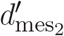 reflects a number of key differences. The fact that temporal bins are grouped into segments (of size 2 in this case) and the corresponding temporal integration of raw neural activations introduce statistical interactions within each group. This changes the structure of the signal (numerator of Eq. (14)) from a linear sum to pairwise combinations within segments, leading to new contributions in the form of the inner products across microscale time bins 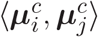. These cross-terms capture how similar signal components across nearby microscale time bins contribute jointly, rather than independently. At the same time, noise (denominator of Eq. (14)) reflects correlated variability within each segment, as captured by the corresponding aggregation terms. Therefore, at both levels, the structure of the d-prime at the mesoscale replaces independent accumulation with localized (within-segment) integration.

Similar to above, we next consider the *coarse mesoscale* representation obtained by integrating neural activations over each half of trial duration. This results in the following closed-form expression of sensitivity index.

#### Theorem 3

(Sensitivity Index at the Coarse Mesoscale). *Let* 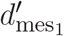 *denote the sensitivity index defined in Eq. (4) at the coarse mesoscale, with similarity statistics* 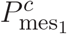 *and* 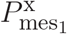 *defined in Eq. (3), trial hypervectors* 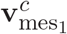 *defined in Eq. (2c), and neural responses following the statistical model in Eq. (5). Assume that Assumption 1 holds. Then*,

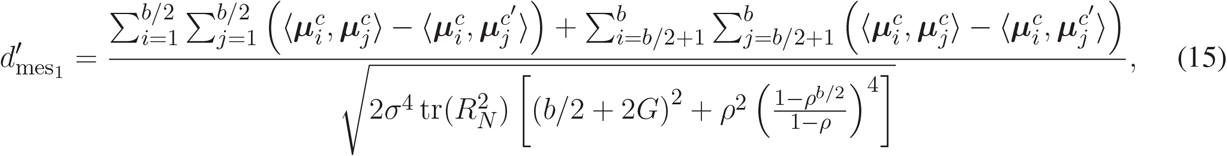

*where*

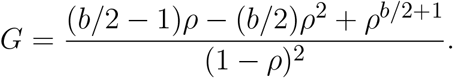

Finally, the following result provides a closed-form expression for d-prime at the macroscale extreme, where neural activity is fully integrated into a single activation vector.

#### Theorem 4

(Sensitivity Index at the Macroscale). *Let* 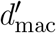 *denote the sensitivity index defined in Eq. (4), with similarity statistics* 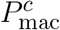 *and* 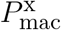 *defined in Eq. (3), trial hypervectors* 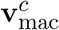 *defined in Eq. (2b), and neural responses following the statistical model in Eq. (5). Assume that Assumption 1 holds. Then*,

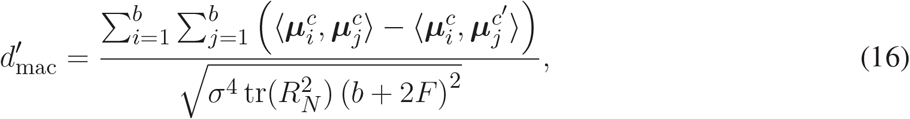

*where*

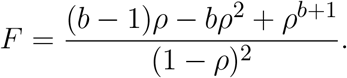

At the macroscale, the aggregation is extended to the entire temporal window, so that all microscale bins are combined into a single representation. Therefore, the signal term in the numerator of Eq. (16) now includes all pairwise interactions across time, accompanied by a respective change in the scaling of the noise term, (*b* + 2*F* )^2^, in the denominator. Notably, this squared form contrasts the linear form in the denominator of Eq. (8) and stems from the fact that temporal integration introduces interactions between all noise components and effectively couples variability across all times. Consequently, noise grows quadratically, making the macroscale highly sensitive to the structure of temporal autocorrelations.

The above results establish how temporal aggregation reshapes both signal and noise contributions across scales. We next leverage these expressions to systematically compare sensitivity across resolutions and determine the parametric regimes in which aggregation enhances or limits discriminability.

#### Numerical Validation of Closed-Form Expressions for Sensitivity Indices

Before analyzing the obtained closed-form expressions for *d*^′^ at various temporal scales, we further validated them against empirical (sample-based) estimates obtained from synthetic neural activity vectors generated according to our statistical model. While the provided analytical proofs are in principle sufficient to guarantee the accuracy of the obtained expressions, this section serves as an additional layer of validation of their correctness.

For each parameter combination (*ρ, β*), two conditions were simulated, each consisting of 60 trials of neural activity recorded over *b* = 20 microscale temporal bins and *N* = 50 neurons. Neural activity vectors were generated according to the model in Eq. (5) with *σ* = *ν* = *k* = 1 and *R*_*N*_ = *I*_*N*_, where condition-dependent mean activity 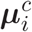 is added to normalized variability 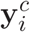. To generate noise vectors with the prescribed temporal autocorrelation structured governed by the decay rate *ρ*, we let

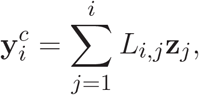

where *L*_*i,j*_ is the (*i, j*)-th element of the *b* × *b* temporal Cholesky factorization *L* of **Σ**_*t*_ (i.e. **Σ**_*t*_ = *LL*^*T*^ ), and **z**_*j*_ ∼ *N*(0, *I*_*N*_ ) represents an independent standard normal spatial noise vector of dimension *N* × 1 drawn at time bin *j*. Similarly, the mean activity vectors 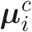 were generated using the Cholesky factor of temporally decaying signal structure in Eq. (7b), controlled by the decay rate *β*. These were further ensured to be embedded in distinct subspaces for the two conditions, ensuring separability in the mean activity patterns while maintaining identical noise statistics. Trial vectors at the micro-, fine meso-, coarse meso-, and macroscale temporal resolutions were then obtained from these neural activity vectors using the corresponding temporal integration in Eq. (1). For each simulated dataset, empirical decodability was computed at each temporal scales using inner-product similarities between trial vectors generated by the synthetic neural activity vectors. Within-class similarities were constructed from all distinct pairs of trials belonging to the same condition, while between-class similarities were constructed from all pairs of trials across the two conditions. These distributions were then substituted directly into Eq. (4) to obtain the empirical *d*^′^ and compared against the analytical expressions from Theorems 1-4.

As shown in Figure 2, the theoretical predictions closely match the empirical estimates across the entire (*ρ, β*) parameter space for all temporal scales. Also notable is the more variable and “noisy” nature of sample estimates, which only recover the ground truth in the limit of infinite samples. Therefore, in addition to validating the accuracy of the obtained analytical expressions, this comparison further highlights the value of the latter, which will become clearer as we subsequently analyze the subtle and intricate optimality transition boundaries among scales.

**Figure 2.**
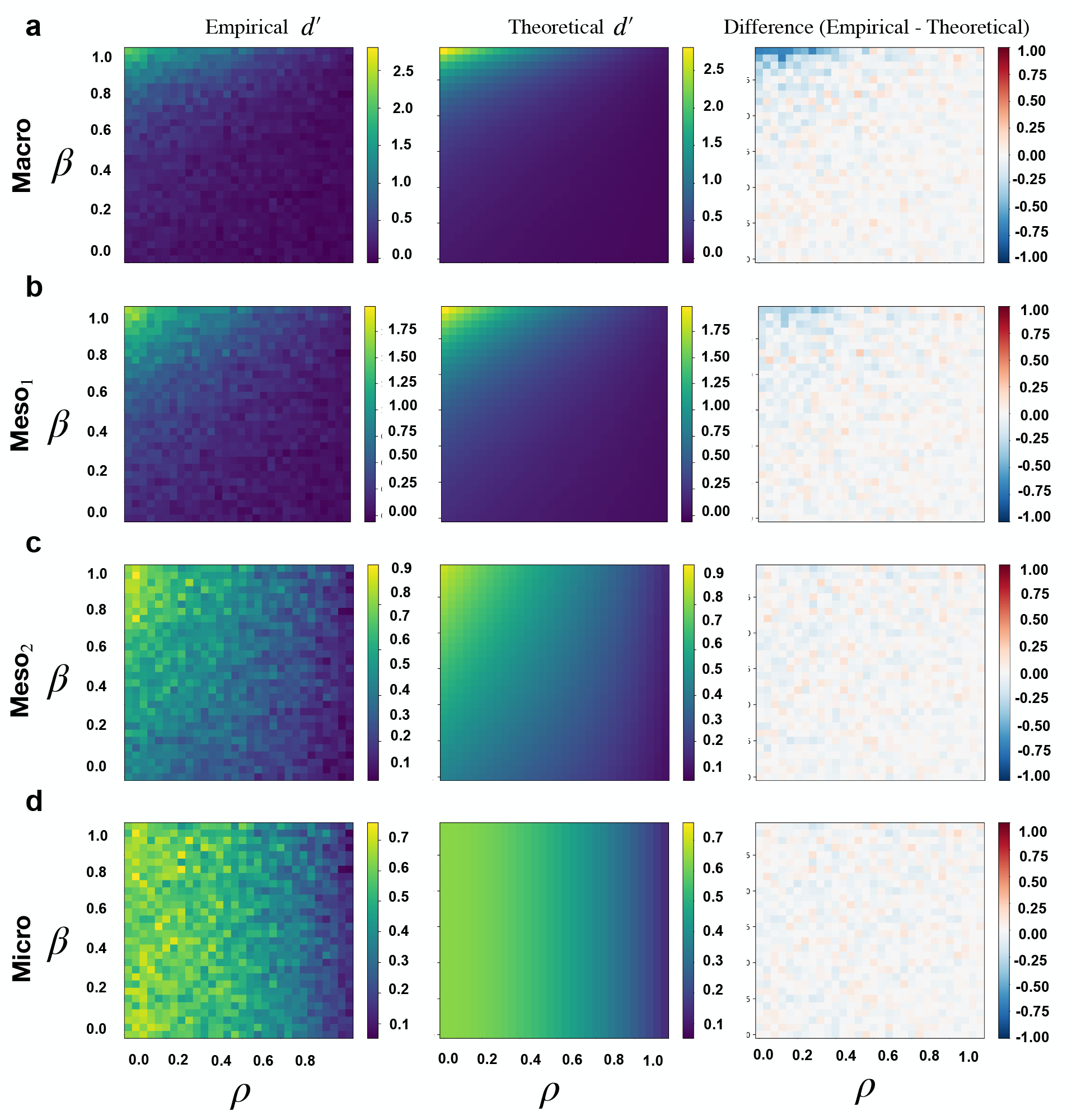
Empirical validation of theoretical expressions for sensitivity index across temporal scales. **(a)** Heatmaps of empirical and theoretical sensitivity index *d*^′^ at the macroscale, together with the difference between the two, shown over the full range of (*ρ, β*) parameter space. **(b–d)** Corresponding heatmaps and differences for the coarse mesoscale, fine mesoscale, and microscale representations, respectively.

### Comparison Across Temporal Scales

The analytical expressions provided in Theorems 1-4 allow us to analyze how d-prime varies across temporal scales for different values of signal and noise autocorrelations. Before analyzing the general cases corresponding to arbitrary values of *β* and *ρ*, we begin with two limiting regimes that more clearly illustrate the contrasting roles of signal and noise autocorrelations. Across all cases we assume the signal to be uncorrelated across conditions, namely, that 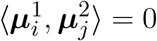 for all *i, j*.

#### Case 1: uncorrelated signal and noise across time

We begin with the extreme scenario where neither signal nor noise are correlated over time, namely,

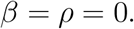

Substituting these into the expressions for the sensitivity index *d*^′^ at the four scales (Eq. (8), Eq. (15), Eq. (14), and Eq. (16)), yields

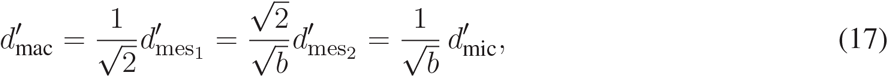

showing that both mesoscale and macroscale representations lead to a *reduction* in the sensitivity index relative to the microscale. In other words, in the absence of signal and noise correlations, averaging degrades decodability and raw neural activations at full resolution provide the highest amount of information about task conditions.

This observation can be understood intuitively by viewing each microscale time bin as carrying an independent piece of information about the condition in the absence of temporal structure. In this case, temporal integration does not introduce new information; instead, it aggregates independent signal contributions while smoothing out independent noise fluctuations, leading to reduced decodability.

Analytically the inner-product statistics are affected by temporal integration in the absence of autocorrelations. At the microscale, the trial hypervector 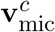 preserves all constituent information, and the inner products in Eq. (3) for microscale, decompose into a sum of independent bin-wise contributions. Consequently, the difference between same-class and different-class inner products, i.e., 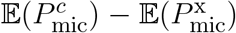, accumulates linearly across microscale bins, reflecting independent signal contributions at all microscale time bins. In contrast, at the meso- and macroscales, multiple microscale temporal bins are combined into a reduced number of effective components. Since the signal has no temporal structure to exploit, this integration does not recover additional information and instead reduces the number of independent signal contributions represented in the inner products, thereby decreasing the signal term in the numerator of Eq. (4). On the other hand, the variance terms in the denominator of Eq. (4) scale linearly with *b* due to the additive accumulation of independent noises, leading to the deterioration of the signal-to-noise ratio encoded by d-prime under averaging.

#### Case 2: fully correlated signal but uncorrelated noise across time

Next, consider the other extreme scenario where noise components still remain uncorrelated but signal remains perfectly correlated over time, namely,

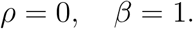

Note that when *β* = 1, the two correlation structures in Eq. (7) coincide. It is straightforward to see that in this case signal must be constant, i.e.,

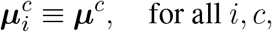

Intuitively, in this case every microscale time bin carries the same signal content, and thus repeated observations across time are redundant in terms of signal but provide repeated opportunities for noise averaging. Temporal integration therefore does not aggregate new signal information; instead, it reinforces the same signal coherently while averaging out independent noise fluctuations, leading to improved decodability. Theoretically, we obtain

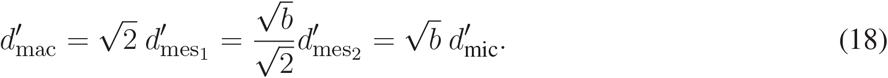

These relations show that, in this case, temporal integration leads to an increase in the sensitivity index, with the macroscale achieving the largest gain. This behavior arises from how temporal integration interacts with the inner-product structure in the presence of aligned signal. In contrast to Case 1, integration at the meso and macro scales now combines microscale bins that carry identical signal components.

Therefore, signal contributions reinforce coherently across time, leading to a superlinear increase in the numerator of Eq. (4) through pairwise interactions between time points. At the same time, since noise remains uncorrelated across time, the variance terms in the denominator of Eq. (4) still scale only through independent contributions. Consequently, noise grows more slowly compared to the coherently reinforced signal.

Together, Cases 1 and 2 represent two extreme scenarios with optimal condition decodability attained at, correspondingly, extreme scales. We next move to more general cases where signal and noise autocorrelations can vary parametrically according to Eq. (6) and Eq. (7).

#### Case 3: persistent signal and decaying noise autocorrelations over time

We next consider the general regime in which noise autocorrelations decay with an arbitrary rate *ρ* according to Eq. (6), while signal correlations remain persistent at an arbitrary value *β* following Eq. (7a). Assume, further, that the magnitude of signal is approximately constant, namely, that

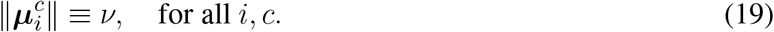

Substituting these into the expressions for *d*^′^ in Theorems 1-4 yields

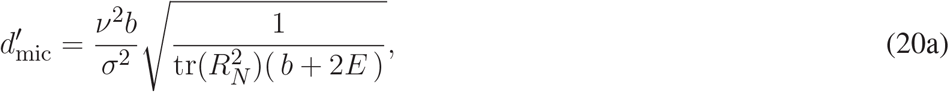

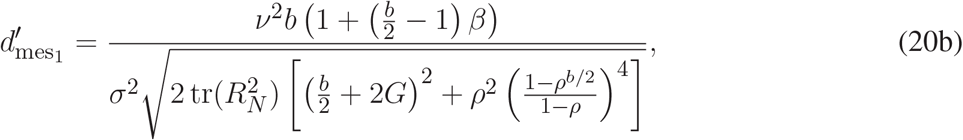

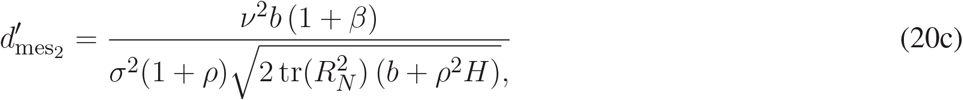

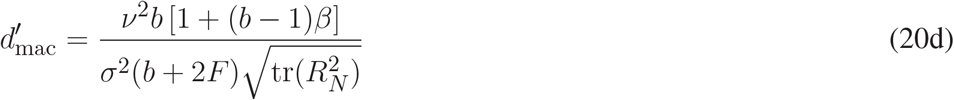

where the variables *E, F*, and *G* are given in Theorems 1-4 and further depend on *ρ*. Unlike the previous two cases, the sensitivity index at neither scale is universally bigger than another. Instead, the relationship between each pair depends on the correlation parameters *β* and *ρ*. In particular, for *s, s*^′^ ∈ *S* it can be shown that

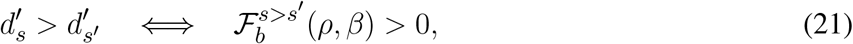

where closed-form expressions for the functions 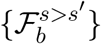 (which depend, notably, on the number *b* of time points) are provided in Supplementary Table S1. These inequalities provide a complete characterization of the parameter regimes governing the relative optimality of each scale.

Figure 3a demonstrates the precise boundary in the *ρ*-*β* plane where each comparison flips its direction, as well as a global illustration of which scale attains the greatest d-prime for each (*ρ, β*) combination. Notably, across all pairwise comparisons, the marked boundaries are monotonically increasing, giving rise to two regions in the *ρ*-*β* plane: a top-left region with relatively large *β* and small *ρ*, where more temporal integration is beneficial, and a bottom-right region with relatively small *β* and large *ρ*, where finer scales have greater d-prime. This behavior can be understood through the balance between signal and noise integration across temporal scales. When signal autocorrelation (*β*) is large and noise autocorrelation (*ρ*) is small, temporal integration largely preserves the shared signal but weakens (cancels-out) noise across time, resulting in improved signal to noise ratio and better decodability. In contrast, when noise autocorrelation is larger than signal autocorrelation, temporal integration has the opposite effect, making finer temporal scales more decodable.

**Figure 3.**
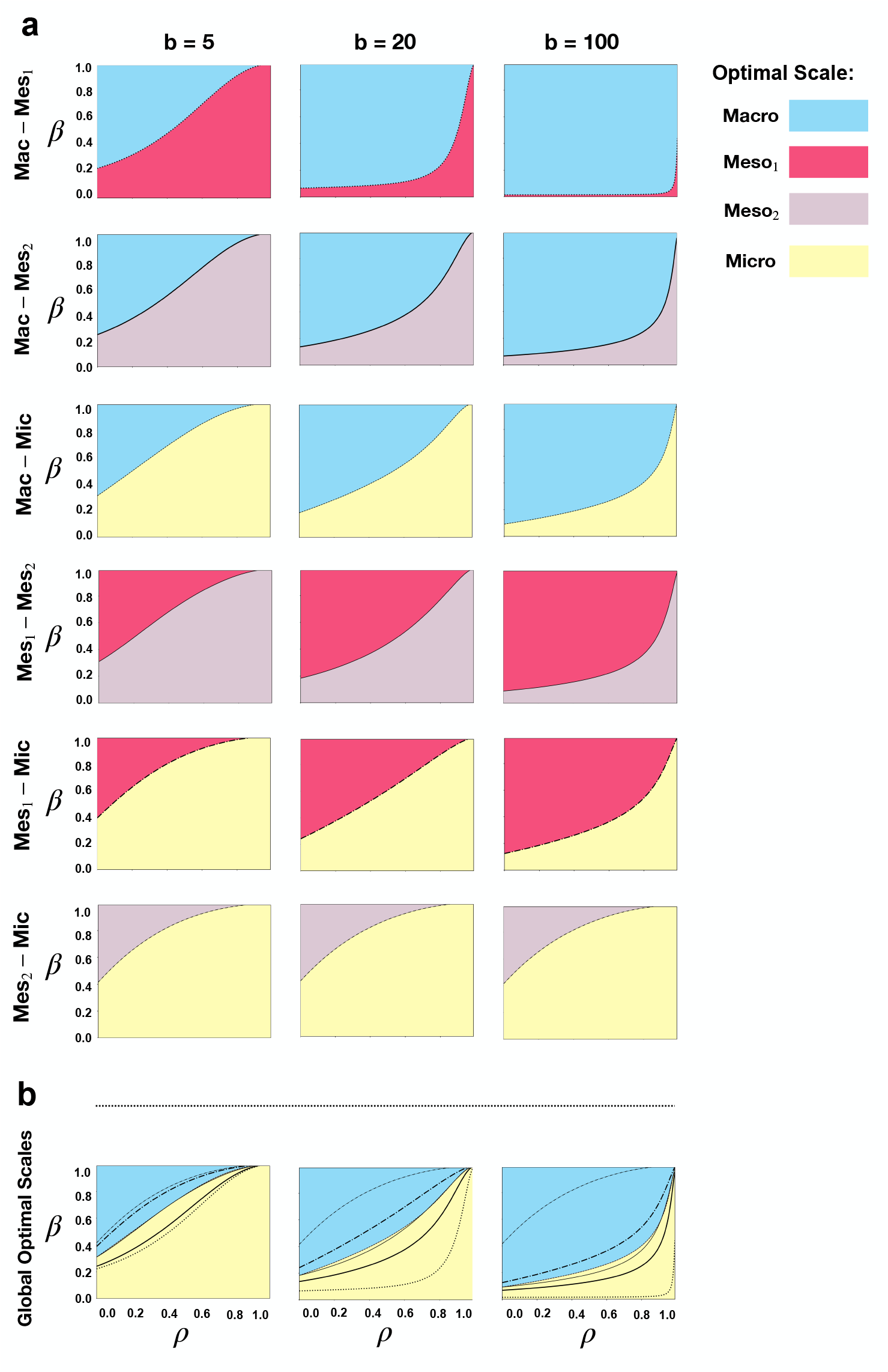
Optimal temporal resolution for information representation under persistent signal auto-correlations. **(a)** Regional depictions of the pairwise comparisons under persistent signal and decaying noise autocorrelations and varying number *b* of microscale time bins. Pairwise comparisons are shown between micro-, fine meso-, coarse meso-, and macroscale (cf. Eq. (21)). The boundary curves indicates the correlation values where 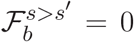 for all *s, s*^′^ ∈ *S*, and the colors denote the scale with greater sensitivity index (d-prime). **(b)** The overlaid comparison of all four scales, summarized from (a). While all six boundaries are shown for ease of comparison, colors denote the optimal scale at the respective (*ρ, β*) combinations.

In addition to the correlation parameters *β* and *ρ*, the relationship between scales depends notably on the number *b* of time points. In Figure 3a, as *b* increases (left to right). The boundary at which the inequality switches (i.e.,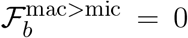) progressively shifts toward the bottom right, implying that temporal integration becomes more beneficial *for the same* (*ρ, β*) as the number of time points increases. This behavior occurs because the signal persists over time (cf. Eq. (7a)), allowing its effects to accumulate with integration over larger number of time points. Meanwhile, as the noise autocorrelations decay over time, integration continues to have stronger weakening effects on noise over larger windows. A similar trend is observed across all pairwise comparisons reinforcing the presence of a strong and robust benefit in temporal integration over larger windows in the presence of persistent signal correlation.

When considering all four scales together, the optimal resolution falls—depending on the relative strength of signal and noise autocorrelations—either at the micro or the macro scale, but never at either of the mesoscales (see Figure 3b). This exclusive behavior shows the strength of the dichotomous tradeoff mentioned earlier, where the existence of a persistent signal correlation results in a marked difference between the effect of integration on signal and that on noise. As a result, the optimal *d*^′^ is driven toward the two extremes, never favoring an intermediate resolution. This behavior is in contrast, in particular, to our empirical findings of mesoscale optimality by Samiei et al. (2024), and motivated us to extend our framework to the more general case of Eq. (7b), as discussed next.

#### Case 4: decaying signal and noise autocorrelations over time

The preceding analysis illustrated the effect of temporal integration on neural representations within extreme regimes in which signal and noise autocorrelations were absent or remained constant across time. While these limiting cases and their universal optimality of extreme scales provide significant insight into the interplay between temporal integration and temporal signal/noise correlations, we next analyze the likely more realistic regime in which both signal and noise autocorrelations decay with temporal distance following Eq. (6) and Eq. (7b). Substituting these into Eq. (8), Eq. (15), Eq. (14) and Eq. (16), we obtain expressions analogous to Eq. (20), with the key difference that signal contributions now involve geometric sums of the form ∑_*i,j*_ *β*^|*j*−*i*|^.

Interestingly, however, since the microscale sensitivity index in Eq. (8) does not involve average neural activity interaction terms across temporal bins, it remains unaffected by the introduction of such temporal correlations and 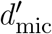 is still given by Eq. (20a). In contrast, the meso- and macroscale sensitivity indices are all influenced by the presence of decaying temporal correlations in signal, and can be shown to be given

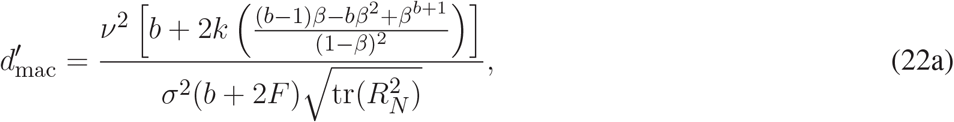

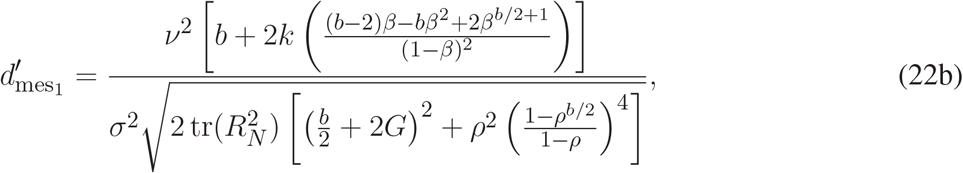

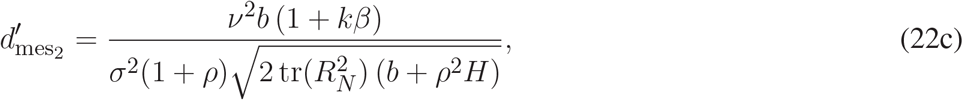

where *F, G* and *H* are given in Theorems 2-4. Similar to Eq. (21), for *k* = 1, pairwise comparisons between these expressions can be simplified to

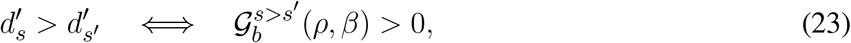

where the functions 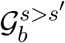 are given in Supplementary Table S2.

Figure 4a demonstrates the regions in the *ρ*-*β* plane where each inequality in Eq. (23) is (or is not) valid for the case where *k* = 1. Similar to Figure 3a, all six pairwise boundaries (i.e 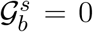) remain monotonically increasing, dividing the parameter space into high *ρ*/low *β* and low *ρ*/high *β* regions where the finer and coarser scales achieve greater decodability, respectively. In contrast to Figure 3a, however, increasing the number of microscale temporal bins *b* now has the effect of shifting all pairwise boundaries upwards, making d-prime greater at finer scales. This indicates that temporal integration is more likely to become ‘excessive’ when signal correlations decay over time, an effect that is consistent with the larger signal cancellation that takes place as a result of temporal averaging under decaying autocorrelations.

**Figure 4.**
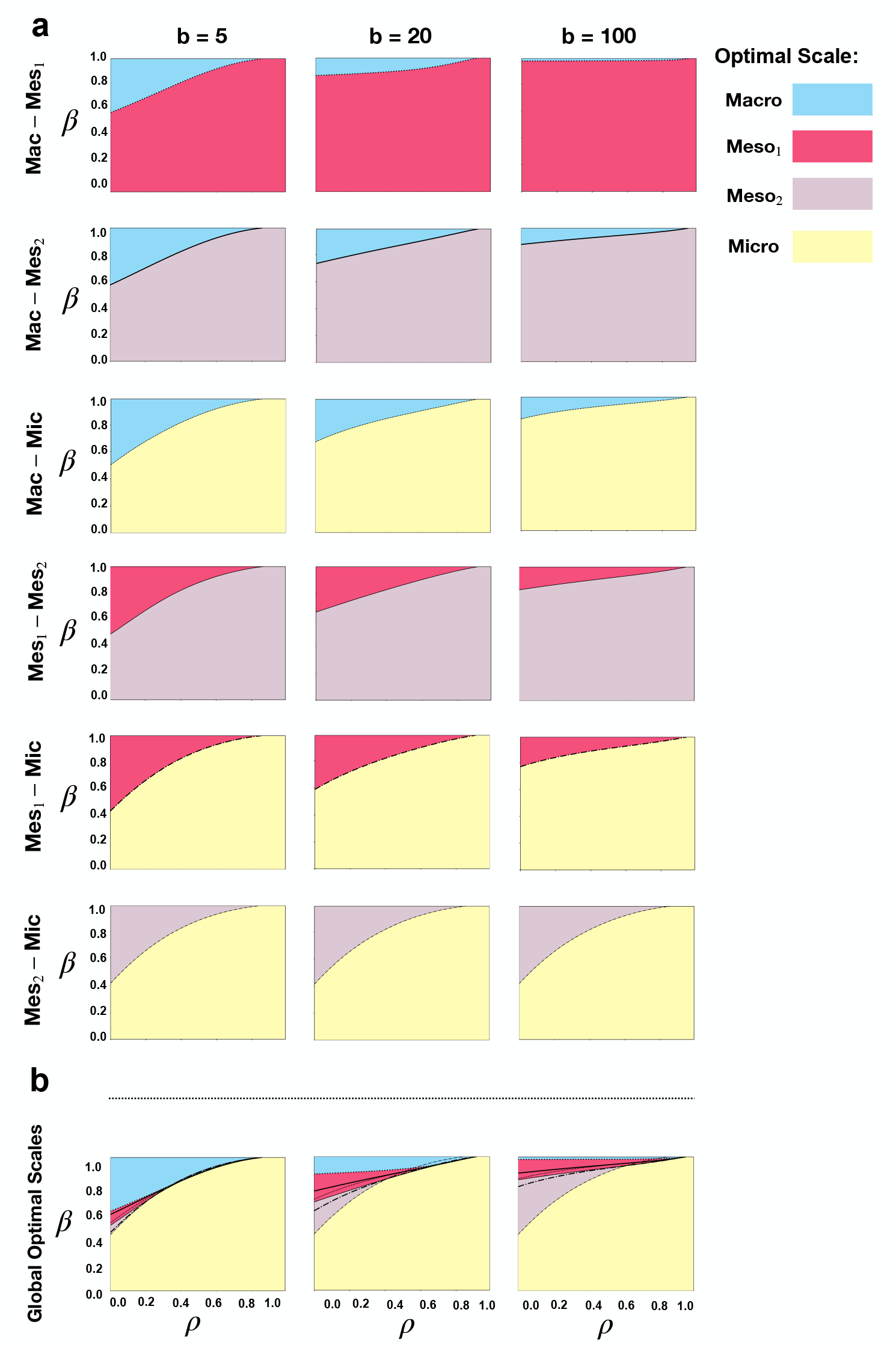
Optimal temporal resolution for information representation under decaying signal auto-correlations. **(a)** Similar to 3(a) but showing the pairwise comparisons of optimal regions for decaying noise and signal autocorrelations given by Eq. (23). Here boundary curves are given by 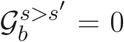. **(b)** Overlaid summary of all six comparisons in (a). Details parallel those in 3(b), but now we observe all four scales to be optimal for a corresponding region in the *ρ*-*β* plane.

Most notably, unlike the case with persistent signal correlations, both the finer and the coarse mesoscale now emerge as the optimal resolution for suitable *ρ, β* in the summarized comparison across all four scales as depicted in Figure 4b. While the microscale dominates in most parameter regimes and the macroscale optimality region shrinks with increasing *b*, the mesoscale optimality regions persists across all *b*. The latter further grows with increasing *b*, particularly in biologically realistic parameter regimes characterized by slow decay in signal autocorrelations and fast decay in noise autocorrelations. This finding is remarkable given our recent observation in (Samiei et al., 2024) of the optimality of the mesoscale in real neural data. Moreover, mesoscale optimality also persists for different values of *k*, as shown in Supplementary Figure S2, although the balance generally tilts towards the microscale with decreasing *k*, as expected. Overall, our results demonstrate that introducing decaying signal correlations fundamentally reshapes the optimality landscape, giving rise to a robust intermediate regime under biologically realistic signal-to-noise conditions, whereby mesoscale averaging provides the optimal balance between capturing coherent information-bearing signal while averaging over task-irrelevant fluctuations.

## DISCUSSION

### Summary

In this work we developed a rigorous theoretical framework showing that the optimal temporal scale of information representation in neural populations is determined by the balance between microscale signal autocorrelations, microscale noise autocorrelations, and the way both decay across time. Through an explicit analysis of the decodability of task conditions from temporally integrated population activity at different scales, we showed that different autocorrelation regimes give rise to different optimal scales of representation. Most notably, in the extreme cases where signal and noise correlations are either absent or persistent throughout the trial duration (Cases 1-3, see Results), the optimal resolution falls at one of the two extremes: macroscale (microscale) if signal autocorrelations are significantly larger (smaller) than noise autocorrelations. On the other hand, if both signal and noise autocorrelations decay with time (Case 4), mesoscale representations also become optimal for a robust and realistic range of parameters: where signal autocorrelations decay slowly while noise autocorrelations decay significantly faster. This effect was observed robustly for all values of trial duration (*b*) and initial signal autocorrelation decay (*k*), with increasing *b* and decreasing *k* both tilting the overall balance towards finer scales.

### The dual effects of averaging

Our results clearly demonstrate a fundamental phenomenon where temporal integration is a double-edged operation for neural representations: it can improve task-relevant information representation by averaging out noise across time, but it can also weaken such information by canceling signal components that are only weakly correlated across temporal bins. This effect can be seen most strongly when both signal and noise autocorrelations decay with temporal separation (Case 4, see Results). In this case temporal integration gives rise to a genuine trade-off: modest integration enhances signal-to-noise ratio and decodability, whereas excessive integration disrupts representational structure and lowers decodability. In this biologically more realistic regime, mesoscale representations emerge as optimal.

These results further provide a theoretical explanation for our recent empirical observations of mesoscale optimality in neural systems (Samiei et al., 2024). Rather than being an intrinsic property of neural coding, mesoscale optimality arises as a consequence of structured temporal correlations. More broadly, our results highlight that temporal averaging is not merely a preprocessing step, but a fundamental operation that reshapes neural information in a structured and predictable manner.

### Role of signal–noise interactions

Our main analytical results are derived under the simplifying Assumption 1 that signal and noise components are uncorrelated. This assumption allows the sensitivity index to decompose explicitly into signal and noise contributions and therefore leads to tractable closed-form expressions. Under this setting, the sensitivity index effectively behaves as a signal-to-noise ratio, where signal accumulation and noise variance (numerator and denominator of d-prime, respectively) can be analyzed independently across temporal scales.

Nevertheless, our general approach and theoretical framework can be extended to the general setting where signal and noise components are not necessarily independent. An example of such extension is provided in Supplementary Note 2 for the case of two time points (*b* = 2) and independence among neurons (*R*_*N*_ = *I*_*N*_ ). Interestingly, while relaxing Assumption 1 introduces signal–noise interaction terms which make closed-form expressions considerably more cumbersome, it does not qualitatively change the model’s behavior. Within this setting, the sensitivity index depends explicitly on both signal magnitude (*ν*^2^) and noise magnitude (*σ*^2^). When noise dominates (*σ*^2^ ≫ *ν*^2^), the behavior closely matches the results obtained under Assumption 1. In contrast, when signal dominates (*ν*^2^ ≫ *σ*^2^), temporal averaging becomes uniformly beneficial. For intermediate cases (e.g., *ν* = *σ* = 1, Supplementary Figure S1a), the general model shows a stronger preference for averaging compared to the simplified case, reflecting the contribution of signal–noise interaction terms. Although the optimality curves for averaging and non-averaging still intersect, the resulting feasible regions are slightly shifted relative to the setting with no signal-noise interactions (Supplementary Figure S1).

### Implications for neural data analysis

Our results have direct implications for how neural data should be analyzed and interpreted. Many standard preprocessing techniques—such as temporal binning, smoothing, and dimensionality reduction—implicitly perform temporal averaging. Our framework shows that the effectiveness of such operations in ‘cleaning the signal’ and extracting meaningful task-relevant dynamics from task-irrelevant noise depends critically on the temporal correlation structure of the data. In particular, in task conditions and brain structures where signal correlations persist much longer than noise correlations (e.g., under stationary tasks or within higher-order association cortices), averaging improves signal to noise ratio up to significantly coarser scales compared to task conditions and brain structures where signal correlations decay relatively fast (e.g., under highly dynamic tasks or within early sensory structures). More broadly, our result suggests that optimal preprocessing pipelines should not rely on fixed temporal scales, but should instead adapt to the underlying signal and noise statistics of the data, and the theoretical conditions derived in this work provide a principled basis for such adaptations.

### Limitations and future directions

For analytical tractability, we considered a restricted set of assumptions. First, we assumed signal and noise autocorrelations that were either zero, constant, or exponentially decaying. While these capture key regimes of temporal dependence, real neural activity may exhibit more complex correlation structures. Second, we assumed noise to be independent across neurons and focused solely on statistical dependence over time. Extending the framework to incorporate spatial correlations and network structure would provide a more complete picture of multiscale neural representations and remains an important direction for future work.

A further natural extension of this framework is to examine whether and to what extent the predicted optimal scales vary across recording modalities. In our earlier empirical work using single-unit spiking activity recorded via NeuroPixel probes by Samiei et al. (2024), we observed specific spatial and temporal mesoscales to be optimal for decoding visual stimulus identity. The present theory suggests that those optimal resolutions are not universal, but instead depends on the underlying structure of signal and noise correlations that may differ across data modalities. Testing this prediction (both directly and via a combination of results developed here and quantification of correlation decay rates) in diverse data modalities can shed light on interesting interplays between optimal spatiotemporal resolutions, data modality, and temporal dynamics.

## METHODS

Detailed proofs of our analytical results (Theorems 1-4), constituting the core part of our methodology, are provided in Supplementary Note 1. In this section we provided additional details about the computational methods we used to interpret the results of these Theorems and compare decodability across scales.

### Analytical derivation of pairwise comparison functions across scales

To characterize the relative optimality of different temporal scales, we formulated pairwise comparisons in terms of inequality conditions between the corresponding sensitivity indices. Specifically, we defined optimality condition functions in Eq. (21) and Eq. (23), with closed-form expressions given in Supplementary Tables S1 and S2, respectively. These functions are obtained by substituting the closed-form expressions of *d*^′^ into the pairwise differences and simplifying the resulting expressions. To ensure the validity of these inequalities, we analyzed the dominant terms of each expression and examined their dependence on the parameters *ρ* and *β*. In particular, we verified that the leading-order contributions remain positive across the parameter regimes considered. Formally, we computed the following partial derivatives and verified that they satisfy

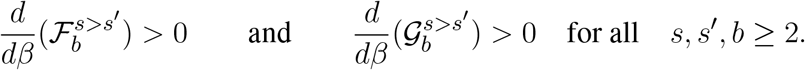

This confirms the monotonic behavior of the expressions and therefore ensures that the sign of each inequality is preserved within the relevant domain.

### Numerical evaluation and visualization of optimality regions

To visualize the analytically derived optimality conditions, we evaluate the functions 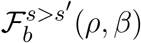 and 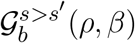 (cf. Supplementary Tables S1 and S2) numerically over a uniform grid defined by *ρ* ∈ [0, 0.99], *β* ∈ [0, 0.99], and a discretization of 300 × 300 points (no numerical root-finding). For each grid point, the sign of the corresponding function determines the optimal regime, with the convention that

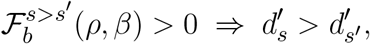

and vice versa. The resulting regions were then visualized using filled contour plots, where Boolean masks identify parameter regions corresponding to each optimal regime. The boundaries between regimes are given by the zero level sets

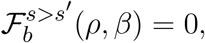

which were approximated numerically as contour lines on the discretized grid. To avoid numerical instabilities associated with singularities in the analytical expressions, the parameter range is restricted to *ρ <* 1. The same process was repeated for the comparison functions 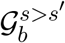 visualized in Figure 4.

## STATEMENTS

### Author contributions

EN designed and supervised the study; HFA derived the analytical results and performed the numerical analyses; TS assisted with numerical analyses and interpretations related to hyper-dimensional computing; HFA drafted and all authors edited the manuscript.

