## Supplementary Material for "On the Optimal Temporal Resolution for Information Representation in Neural Activity: A Theoretical Analysis"

### SUPPLEMENTARY NOTE 1: ANALYTICAL PROOFS

#### *Sensitivity Index at the Microscale*

PROOF OF THEOREM 1. Let $\mathbf{v}^c_{\text{mic}}$ denote the trial vector at microscale resolution defined as:

$$\mathbf{v}^c_{\text{mic}} = \sum_{i=1}^{b} \mathbf{S}\left(\boldsymbol{\mu}^c_i + \sigma \mathbf{y}^c_i\right) \odot \boldsymbol{t}_i.$$

Let $T_i = \text{diag}(\boldsymbol{t}_i) \in \mathbb{R}^{D \times D}$. Then we can equivalently write

$$\mathbf{v}^c_{\text{mic}} = \sum_{i=1}^{b} T_i \mathbf{S}\left(\boldsymbol{\mu}^c_i + \sigma \mathbf{y}^c_i\right).$$

Now to define within-class similarities for microscale trial hypervectors, we have,

$$\begin{aligned} P^c_{\text{mic}} &= \left(\sum_{i=1}^{b} T_i \mathbf{S}(\boldsymbol{\mu}^c_i + \sigma \mathbf{y}^c_i)\right)^{\top} \left(\sum_{k=1}^{b} T_k \mathbf{S}(\boldsymbol{\mu}^c_k + \sigma \hat{\mathbf{y}}^c_k)\right) \\ &= \sum_{i=1}^{b} \sum_{k=1}^{b} (\boldsymbol{\mu}^c_i + \sigma \mathbf{y}^c_i)^{\top} \mathbf{S}^{\top} T_i^{\top} T_k \mathbf{S}(\boldsymbol{\mu}^c_k + \sigma \hat{\mathbf{y}}^c_k). \end{aligned}$$

Since $\boldsymbol{t}_i$ are orthogonal, we have $T_i^{\top} T_k = 0$ for $i \neq k$ and $T_i^{\top} T_i = I$ . Also $\mathbf{S}^{\top}\mathbf{S} \approx \gamma I_N$. Hence,

$$P^c_{\text{mic}} = \gamma \sum_{i=1}^{b} (\boldsymbol{\mu}^c_i + \sigma \mathbf{y}^c_i)^{\top} (\boldsymbol{\mu}^c_i + \sigma \hat{\mathbf{y}}^c_i).$$

Expanding each term,

$$(\boldsymbol{\mu}^c_i + \sigma \mathbf{y}^c_i)^{\top} (\boldsymbol{\mu}^c_i + \sigma \hat{\mathbf{y}}^c_i) = \|\boldsymbol{\mu}^c_i\|^2 + \sigma \boldsymbol{\mu}^{c\top}_i \hat{\mathbf{y}}^c_i + \sigma \mathbf{y}^{c\top}_i \boldsymbol{\mu}^c_i + \sigma^2 \mathbf{y}^{c\top}_i \hat{\mathbf{y}}^c_i. \tag{S1}$$

Taking expectation and using $\mathbb{E}[\mathbf{y}^c_i] = 0$ together with assumption 1 and independence across trials,

$$\mathbb{E}[P^c_{\text{mic}}] = \gamma \sum_{i=1}^{b} \|\boldsymbol{\mu}^c_i\|^2. \tag{S2}$$

Similarly, for between-class i.e $c \neq c'$, we have,

$$P^{\text{x}}_{\text{mic}} = \gamma \sum_{i=1}^{b} (\boldsymbol{\mu}^c_i + \sigma \mathbf{y}^c_i)^{\top} (\boldsymbol{\mu}^{c'}_i + \sigma \hat{\mathbf{y}}^{c'}_i).$$

Taking expectation,

$$\mathbb{E}[P^{\mathrm{x}}_{\mathrm{mic}}] = \gamma \sum_{i=1}^{b} \langle \boldsymbol{\mu}_i^c, \boldsymbol{\mu}_i^{c'} \rangle. \tag{S3}$$

We now compute the variance of $P^c_{\mathrm{mic}}$:

$$\mathrm{Var}[P^c_{\mathrm{mic}}] = \gamma^2 \mathrm{Var}\Big[\sum_{i=1}^{b} \|\boldsymbol{\mu}_i^c\|^2 + \sigma \mathbf{y}_i^{c\top} \boldsymbol{\mu}_i^c + \sigma \boldsymbol{\mu}_i^{c\top} \hat{\mathbf{y}}_i^c + \sigma^2 \mathbf{y}_i^{c\top} \hat{\mathbf{y}}_i^c\Big].$$

Since deterministic terms and linear terms vanish under variance, we obtain

$$\mathrm{Var}[P^c_{\mathrm{mic}}] = \gamma^2 \sigma^4 \mathrm{Var}\Big[\sum_{i=1}^{b} \mathbf{y}_i^{c\top} \hat{\mathbf{y}}_i^c\Big].$$

Expanding the variance,

$$\mathrm{Var}[P^c_{\mathrm{mic}}] = \gamma^2 \sigma^4 \mathbb{E}\Big[\sum_{i=1}^{b} (\mathbf{y}_i^{c\top} \hat{\mathbf{y}}_i^c)^2 + 2 \sum_{1 \le i < j \le b} \mathbf{y}_i^{c\top} \hat{\mathbf{y}}_i^c \mathbf{y}_j^{c\top} \hat{\mathbf{y}}_j^c\Big].$$

Using the trace identity $a^\top b\, b^\top a = \mathrm{tr}(aa^\top bb^\top)$,

$$\mathrm{Var}[P^c_{\mathrm{mic}}] = \gamma^2 \sigma^4 \Bigg( \sum_{i=1}^{b} \mathrm{tr}\left(\mathbb{E}[\mathbf{y}_i^c \mathbf{y}_i^{c\top}] \mathbb{E}[\hat{\mathbf{y}}_i^c \hat{\mathbf{y}}_i^{c\top}]\right) + 2 \sum_{1 \le i < j \le b} \mathrm{tr}\left(\mathrm{Cov}(\mathbf{y}_i^c, \mathbf{y}_j^c)\, \mathrm{Cov}(\hat{\mathbf{y}}_i^c, \hat{\mathbf{y}}_j^c)\right) \Bigg).$$

Using $\mathrm{Cov}(\mathbf{y}_i^c, \mathbf{y}_j^c) = \rho^{|j-i|} R_N$, we obtain

$$\mathrm{tr}(R_N R_N) = \mathrm{tr}(R_N^2), \qquad \mathrm{tr}(\rho^{j-i} R_N \cdot \rho^{j-i} R_N) = \rho^{2(j-i)}\, \mathrm{tr}(R_N^2).$$

Thus,

$$\mathrm{Var}[P^c_{\mathrm{mic}}] = \gamma^2 \sigma^4\, \mathrm{tr}(R_N^2) \left( b + 2 \sum_{1 \le i < j \le b} \rho^{2(j-i)} \right). \tag{S4}$$

We now simplify the series in Eq. (S4):

$$\sum_{1 \le i < j \le b} \rho^{2(j-i)} = \sum_{k=1}^{b-1} (b-k) \rho^{2k} \tag{S5a}$$

$$= b \sum_{k=1}^{b-1} \rho^{2k} - \sum_{k=1}^{b-1} k \rho^{2k} \tag{S5b}$$

$$= b\frac{(\rho^2-\rho^{2b})}{1-\rho^2} - \frac{\rho}{2}\frac{d}{d\rho}\sum_{k=0}^{b-1}\rho^{2k} \tag{S5c}$$

$$= b\frac{(\rho^2-\rho^{2b})}{1-\rho^2} - \frac{\rho}{2}\frac{d}{d\rho}\left(\frac{1-\rho^{2b}}{1-\rho^2}\right) \tag{S5d}$$

$$= b\frac{(\rho^2-\rho^{2b})}{1-\rho^2} + \frac{\rho}{2}\left(\frac{(1-\rho^2)(2b\rho^{2b-1})-(2\rho)(1-\rho^{2b})}{(1-\rho^2)^2}\right) \tag{S5e}$$

$$= \frac{(b-1)\rho^2 - b\rho^4 + \rho^{2b+2}}{(1-\rho^2)^2} \tag{S5f}$$

Let us define,

$$\sum_{1\leq i<j\leq b}\rho^{2(j-i)} = \frac{(b-1)\rho^2 - b\rho^4 + \rho^{2b+2}}{(1-\rho^2)^2} = E.$$

Hence after substituting the value of $\sum_{1\leq i<j\leq b}\rho^{2(j-i)}$ in Eq. (S4),

$$\mathrm{Var}[P^c_{\mathrm{mic}}] = \gamma^2\sigma^4\,\mathrm{tr}(R_N^2)(b+2E). \tag{S6}$$

By identical arguments,

$$\mathrm{Var}[P^{\mathrm{x}}_{\mathrm{mic}}] = \gamma^2\sigma^4\,\mathrm{tr}(R_N^2)(b+2E). \tag{S7}$$

Substituting expectation expressions in Eq. (S2) Eq. (S3) and variances in Eq. (S6) Eq. (S7) into the definition of $d'$( Eq. (4) in main text) gives,

$$d'_{\mathrm{mic}} = \frac{\sum_{i=1}^{b}\left(\|\boldsymbol{\mu}_i^c\|^2 - \langle\boldsymbol{\mu}_i^c, \boldsymbol{\mu}_i^{c'}\rangle\right)}{\sqrt{\sigma^4\,\mathrm{tr}(R_N^2)(b+2E)}}. \qquad \square$$

#### *Sensitivity Index at the Fine Mesoscale*

PROOF OF THEOREM 2. Let $\mathbf{v}^c_{\mathrm{mes}_2}$ denote the temporally encoded trial vector at finer mesoscale resolution:

$$\mathbf{v}^c_{\mathrm{mes}_2} = \mathbf{S}\Big(\sum_{i=1}^{2}(\boldsymbol{\mu}_i^c + \sigma\boldsymbol{y}_i^{j,c})\Big)\odot\mathbf{t}_1 + \cdots + \mathbf{S}\Big(\sum_{i=b-1}^{b}(\boldsymbol{\mu}_i^c + \sigma\boldsymbol{y}_i^{j,c})\Big)\odot\mathbf{t}_{b/2}. \tag{S8}$$

proceeding as before with $T$ and $S$ matrices and orthogonality, we have

$$P^c_{\mathrm{mes}_2} = \gamma\Big(\sum_{i=1}^{2}(\boldsymbol{\mu}_i^c + \sigma\mathbf{y}_i^c)\Big)^\top\Big(\sum_{k=1}^{2}(\boldsymbol{\mu}_k^c + \sigma\hat{\mathbf{y}}_k^c)\Big) + \cdots + \gamma\Big(\sum_{i=b-1}^{b}(\boldsymbol{\mu}_i^c + \sigma\mathbf{y}_i^c)\Big)^\top\Big(\sum_{k=b-1}^{b}(\boldsymbol{\mu}_k^c + \sigma\hat{\mathbf{y}}_k^c)\Big)$$

Using $\mathbb{E}[\mathbf{y}_i^c]=0$, within class expectation is,

$$\mathbb{E}[P^c_{\text{mes}_2}] = \gamma \left[ \sum_{i=1}^{b} \|\boldsymbol{\mu}_i^c\|^2 + 2\sum_{i=1}^{b/2} \langle \boldsymbol{\mu}_{2i-1}^c, \boldsymbol{\mu}_{2i}^{c'} \rangle \right] \tag{S9}$$

Similarly, for between-class,

$$\mathbb{E}[P^{\text{x}}_{\text{mes}_2}] = \gamma \left[ \sum_{i=1}^{b} \langle \boldsymbol{\mu}_i^c, \boldsymbol{\mu}_i^{c'} \rangle + \sum_{i=1}^{b/2} \langle \boldsymbol{\mu}_{2i-1}^c, \boldsymbol{\mu}_{2i}^{c'} \rangle + \langle \boldsymbol{\mu}_{2i}^c, \boldsymbol{\mu}_{2i-1}^{c'} \rangle \right]. \tag{S10}$$

Using statistical model properties and assumption 1, in-class variance is,

$$\begin{aligned}
\text{Var}[P^c_{\text{mes}_2}] &= \gamma^2\sigma^4 \text{Var}\Big[ \sum_{i=1}^{b/2} (\mathbf{y}_{2i-1}^c + \mathbf{y}_{2i}^c)^\top (\hat{\mathbf{y}}_{2i-1}^c + \hat{\mathbf{y}}_{2i}^c) \Big] \\
&= \gamma^2\sigma^4 \sum_{i=1}^{b/2} \text{Var}\Big[ (\mathbf{y}_{2i-1}^c + \mathbf{y}_{2i}^c)^\top (\hat{\mathbf{y}}_{2i-1}^c + \hat{\mathbf{y}}_{2i}^c) \Big] \\
&\quad + 2 \sum_{1\le i<k\le b/2} \text{Cov}\Big[ (\mathbf{y}_{2i-1}^c + \mathbf{y}_{2i}^c)^\top (\hat{\mathbf{y}}_{2i-1}^c + \hat{\mathbf{y}}_{2i}^c), (\mathbf{y}_{2k-1}^c + \mathbf{y}_{2k}^c)^\top (\hat{\mathbf{y}}_{2k-1}^c + \hat{\mathbf{y}}_{2k}^c) \Big]
\end{aligned} \tag{S11}$$

The variance term in Eq. (S11) can be simplified as,

$$\text{Var}\big[ (\mathbf{y}_{2i-1}^c + \mathbf{y}_{2i}^c)^\top (\hat{\mathbf{y}}_{2i-1}^c + \hat{\mathbf{y}}_{2i}^c) \big] = \text{tr}(\text{Cov}(\mathbf{y}_{2i-1}^c + \mathbf{y}_{2i}^c)\, \text{Cov}(\hat{\mathbf{y}}_{2i-1}^c + \hat{\mathbf{y}}_{2i}^c)) \tag{S12}$$

where $\text{Cov}(\mathbf{y}_{2i-1}^c + \mathbf{y}_{2i}^c)$ can be expanded as,

$$\begin{aligned}
\text{Cov}(\mathbf{y}_{2i-1}^c + \mathbf{y}_{2i}^c) &= \mathbb{E}\big[ \mathbf{y}_{2i-1}^c \mathbf{y}_{2i-1}^{c\top} + \mathbf{y}_{2i-1}^c \mathbf{y}_{2i}^{c\top} + \mathbf{y}_{2i}^c \mathbf{y}_{2i-1}^{c\top} + \mathbf{y}_{2i}^c \mathbf{y}_{2i}^{c\top} \big] \\
&= \text{Cov}(\mathbf{y}_{2i-1}^c, \mathbf{y}_{2i-1}^c) + \text{Cov}(\mathbf{y}_{2i-1}^c, \mathbf{y}_{2i}^c) + \text{Cov}(\mathbf{y}_{2i}^c, \mathbf{y}_{2i-1}^c) + \text{Cov}(\mathbf{y}_{2i}^c, \mathbf{y}_{2i}^c) \\
&= 2(1+\rho)R_N
\end{aligned}$$

Similarly the later covariance term follows and upon substituting back to Eq. (S12) we have,

$$\text{Var}\big[ (\mathbf{y}_{2i-1}^c + \mathbf{y}_{2i}^c)^\top (\hat{\mathbf{y}}_{2i-1}^c + \hat{\mathbf{y}}_{2i}^c) \big] = 4(1+\rho)^2\, \text{tr}(R_N^2) \tag{S13}$$

Now using the covariance expectation relation and trace operator properties the covariance terms in Eq. (S11) reduces to,

$$\begin{aligned}
&\text{Cov}\Big[ (\mathbf{y}_{2i-1}^c + \mathbf{y}_{2i}^c)^\top (\hat{\mathbf{y}}_{2i-1}^c + \hat{\mathbf{y}}_{2i}^c), (\mathbf{y}_{2k-1}^c + \mathbf{y}_{2k}^c)^\top (\hat{\mathbf{y}}_{2k-1}^c + \hat{\mathbf{y}}_{2k}^c) \Big] \\
&\quad = \text{tr}\left[ \text{Cov}\big( \mathbf{y}_{2i-1}^c + \mathbf{y}_{2i}^c, \mathbf{y}_{2k-1}^c + \mathbf{y}_{2k}^c \big)\, \text{Cov}\big( \hat{\mathbf{y}}_{2i-1}^c + \hat{\mathbf{y}}_{2i}^c, \hat{\mathbf{y}}_{2k-1}^c + \hat{\mathbf{y}}_{2k}^c \big) \right]
\end{aligned}$$

$$
\begin{aligned}
&= \operatorname{tr}\left[(2\rho^{2(k-i)} + \rho^{2(k-i)+1} + \rho^{2(k-i)-1})^2 R_N^2\right] \\
&= (2\rho^{2(k-i)} + \rho^{2(k-i)+1} + \rho^{2(k-i)-1})^2 \operatorname{tr}(R_N^2) \\
&= \rho^{4(k-i)}(2+\rho+\rho^{-1})^2 \operatorname{tr}(R_N^2) \\
&= \rho^{4(k-i)-2}(2\rho+\rho^2+1)^2 \operatorname{tr}(R_N^2)
\end{aligned}
$$

hence,

$$
\operatorname{Cov}\left[(\mathbf{y}_{2i-1}^c + \mathbf{y}_{2i}^c)^\top(\hat{\mathbf{y}}_{2i-1}^c + \hat{\mathbf{y}}_{2i}^c), (\mathbf{y}_{2k-1}^c + \mathbf{y}_{2k}^c)^\top(\hat{\mathbf{y}}_{2k-1}^c + \hat{\mathbf{y}}_{2k}^c)\right] = \rho^{4(k-i)-2}(1+\rho)^4 \operatorname{tr}(R_N^2) \tag{S14}
$$

Hence substituting Eq. (S13) and Eq. (S14) in Eq. (S11) gives,

$$
\operatorname{Var}[P_{\text{mes}_2}^c] = \gamma^2\sigma^4 \left( \operatorname{tr}(R_N^2) \sum_{i=1}^{b/2} 4(1+\rho)^2 + 2\operatorname{tr}(R_N^2) \sum_{1\leq i<k\leq b/2} \rho^{4(k-i)-2}(1+\rho)^4 \right)
$$

$$
\operatorname{Var}[P_{\text{mes}_2}^c] = \gamma^2\sigma^4 \operatorname{tr}(R_N^2) \left( \sum_{i=1}^{b/2} 4(1+\rho)^2 + 2(1+\rho)^4\rho^{-2} \sum_{i=1}^{b/2-1} \sum_{k=i+1}^{b/2} \rho^{4(k-i)} \right) \tag{S15}
$$

The double summation of noise correlation in Eq. (S15) is simplified using finite geometric sum follows,

$$
\begin{aligned}
\sum_{i=1}^{b/2-1} \sum_{k=i+1}^{b/2} \rho^{4(k-i)} &= \sum_{i=1}^{b/2-1} \frac{1}{\rho^{4i}} . \frac{\rho^{4(i+1)} - \rho^{4(b/2+1)}}{1-\rho^4} \\
&= \frac{\rho^4}{1-\rho^4} \sum_{i=1}^{b/2-1} (1-\rho^{2b-4i}) \\
&= \frac{\rho^4}{1-\rho^4}\left[ \left(\frac{b}{2}-1\right) + \frac{(\rho^{2b}-\rho^4)}{1-\rho^4} \right] \\
&= \rho^4 \frac{(b/2-1) - (b/2)\rho^4 + \rho^{2b}}{(1-\rho^4)^2}.
\end{aligned}
$$

Substituting the above expression in Eq. (S15),

$$
\begin{aligned}
\operatorname{Var}[P_{\text{mes}_2}^c] &= \gamma^2\sigma^4 \operatorname{tr}(R_N^2) \left( 2b(1+\rho)^2 + 2(1+\rho)^4\rho^2 \frac{((b/2-1) - (b/2)\rho^4 + \rho^{2b})}{(1-\rho^4)^2} \right) \\
&= 2\gamma^2\sigma^4 \operatorname{tr}(R_N^2)(1+\rho)^2 \left( b + \rho^2 H \right)
\end{aligned} \tag{S16}
$$

where,

$$
H = \frac{(b/2-1) - (b/2)\rho^4 + \rho^{2b}}{(1-\rho)^2(1+\rho^2)^2}.
$$

With similar computation between-class variance is,

$$\mathrm{Var}[P^c_{\mathrm{mes}_2}] = 2\gamma^2\sigma^4\,\mathrm{tr}(R_N^2)(1+\rho)^2\left(b+\rho^2 H\right) \tag{S17}$$

Substituting Eq. (S9) Eq. (S10) Eq. (S16) and Eq. (S17) into Eq. (4) gives the required closed form expression,

$$d'_{\mathrm{mes}_2} = \frac{\sum_{i=1}^{b}\left(\|\boldsymbol{\mu}_i^c\|^2 - \langle \boldsymbol{\mu}_i^c, \boldsymbol{\mu}_j^{c'}\rangle\right) + \sum_{i=1}^{b/2}\left(2\langle \boldsymbol{\mu}_{2i-1}^c, \boldsymbol{\mu}_{2i}^c\rangle - \langle \boldsymbol{\mu}_{2i-1}^c, \boldsymbol{\mu}_{2i}^{c'}\rangle - \langle \boldsymbol{\mu}_{2i}^c, \boldsymbol{\mu}_{2i-1}^{c'}\rangle\right)}{(1+\rho)\sqrt{2\sigma^4\,\mathrm{tr}(R_N^2)\,(b+\rho^2 H)}},$$

### ***Sensitivity Index at the Coarse Mesoscale***

PROOF OF THEOREM 3. Let $\mathbf{v}^c_{\mathrm{mes}_1}$ denote the temporally encoded trial vector at mesoscale temporal resolution, defined as

$$\mathbf{v}^c_{\mathrm{mes}_1} = \mathbf{S}\Big(\sum_{i=1}^{b/2}(\boldsymbol{\mu}_i^c + \sigma \mathbf{y}_i^c)\Big) \odot \mathbf{t}_1 + \mathbf{S}\Big(\sum_{i=b/2+1}^{b}(\boldsymbol{\mu}_i^c + \sigma \mathbf{y}_i^c)\Big) \odot \mathbf{t}_2. \tag{S18}$$

Since the temporal basis vectors $\mathbf{t}_1, \mathbf{t}_2$ are orthogonal, the corresponding diagonal representations $T_1 = \mathrm{diag}(\mathbf{t}_1)$ and $T_1 = \mathrm{diag}(\mathbf{t}_2)$ are also orthogonal, which allows the inner product to decouple across the two halves. Accordingly, the within-class inner product can be written as

$$\begin{aligned} P^c_{\mathrm{mes}_1} = {} & \gamma\Big(\sum_{i=1}^{b/2}(\boldsymbol{\mu}_i^c + \sigma \mathbf{y}_i^c)\Big)^\top \Big(\sum_{i=1}^{b/2}(\boldsymbol{\mu}_i^c + \sigma \hat{\mathbf{y}}_i^c)\Big) \\ & + \gamma\Big(\sum_{i=b/2+1}^{b}(\boldsymbol{\mu}_i^c + \sigma \mathbf{y}_i^c)\Big)^\top \Big(\sum_{i=b/2+1}^{b}(\boldsymbol{\mu}_i^c + \sigma \hat{\mathbf{y}}_i^c)\Big). \end{aligned}$$

Using $\mathbb{E}[\mathbf{y}_i^c] = 0$, the expected within-class inner product becomes

$$\mathbb{E}[P^c_{\mathrm{mes}_1}] = \gamma\left(\sum_{i=1}^{b/2}\sum_{j=1}^{b/2}\langle \boldsymbol{\mu}_i^c, \boldsymbol{\mu}_j^c\rangle + \sum_{i=b/2+1}^{b}\sum_{j=b/2+1}^{b}\langle \boldsymbol{\mu}_i^c, \boldsymbol{\mu}_j^c\rangle\right). \tag{S19}$$

Similarly, the between-class expectation is

$$\mathbb{E}[P^{\mathrm{x}}_{\mathrm{mes}_1}] = \gamma\left(\sum_{i=1}^{b/2}\sum_{j=1}^{b/2}\langle \boldsymbol{\mu}_i^c, \boldsymbol{\mu}_j^{c'}\rangle + \sum_{i=b/2+1}^{b}\sum_{j=b/2+1}^{b}\langle \boldsymbol{\mu}_i^c, \boldsymbol{\mu}_j^{c'}\rangle\right). \tag{S20}$$

We now compute the variance of the within-class inner product:

$$\mathrm{Var}[P^c_{\mathrm{mes}_1}] = \gamma^2\mathrm{Var}\Big[\sum_{i=1}^{b/2}\sum_{j=1}^{b/2}(\boldsymbol{\mu}_i^c + \sigma \mathbf{y}_i^c)^\top(\boldsymbol{\mu}_j^c + \sigma \hat{\mathbf{y}}_j^c)$$

$$+ \sum_{i=b/2+1}^{b} \sum_{j=b/2+1}^{b} (\boldsymbol{\mu}_i^c + \sigma \mathbf{y}_i^c)^\top (\boldsymbol{\mu}_k^c + \sigma \hat{\mathbf{y}}_j^c)\Big].$$

Since signal-noise cross terms vanish and $||\boldsymbol{\mu}||^2$ is deterministic, this reduces to

$$\begin{aligned}
\mathrm{Var}[P^c_{\mathrm{mes}_1}] = \gamma^2\sigma^4 \Big( & \mathrm{Var}\Big[\Big(\sum_{i=1}^{b/2} \mathbf{y}_i^c\Big)^\top \Big(\sum_{i=1}^{b/2} \hat{\mathbf{y}}_i^c\Big)\Big] \\
& + \mathrm{Var}\Big[\Big(\sum_{i=b/2+1}^{b} \mathbf{y}_i^c\Big)^\top \Big(\sum_{i=b/2+1}^{b} \hat{\mathbf{y}}_i^c\Big)\Big] \\
& + 2\ \mathrm{Cov}\Big[\Big(\sum_{i=1}^{b/2} \mathbf{y}_i^c\Big)^\top \Big(\sum_{i=1}^{b/2} \hat{\mathbf{y}}_i^c\Big), \Big(\sum_{i=b/2+1}^{b} \mathbf{y}_i^c\Big)^\top \Big(\sum_{i=b/2+1}^{b} \hat{\mathbf{y}}_i^c\Big)\Big]\Big).
\end{aligned} \tag{S21}$$

We evaluate each term in Eq. (S21) separately. First,

$$\mathrm{Var}\Big[\Big(\sum_{i=1}^{b/2} \mathbf{y}_i^c\Big)^\top \Big(\sum_{i=1}^{b/2} \hat{\mathbf{y}}_i^c\Big)\Big] = \mathrm{tr}\left(\mathrm{Cov}\Big[\sum_{i=1}^{b/2} \mathbf{y}_i^c\Big]\,\mathrm{Cov}\Big[\sum_{i=1}^{b/2} \hat{\mathbf{y}}_i^c\Big]\right). \tag{S22}$$

Using the covariance expectation relation and the model's correlation structure,

$$\begin{aligned}
\mathrm{Cov}\Big[\sum_{i=1}^{b/2} \mathbf{y}_i^c\Big] &= \mathbb{E}\left[\left(\sum_{i=1}^{b/2} \mathbf{y}_i^c\right)\left(\sum_{j=1}^{b/2} \mathbf{y}_j^c\right)^\top\right] \\
&= \sum_{i=1}^{b/2}\sum_{j=1}^{b/2} \mathbb{E}[\mathbf{y}_i^c \mathbf{y}_j^{c\top}] \\
&= \sum_{i=1}^{b/2}\sum_{j=1}^{b/2} \mathrm{Cov}[\mathbf{y}_i^c, \mathbf{y}_j^c] \\
&= \frac{b}{2} R_N + 2 \sum_{1\le i<j\le b/2} \rho^{j-i} R_N.
\end{aligned}$$

Substituting into Eq. (S22) gives

$$\mathrm{Var}\Big[\Big(\sum_{i=1}^{b/2} \mathbf{y}_i^c\Big)^\top \Big(\sum_{i=1}^{b/2} \hat{\mathbf{y}}_i^c\Big)\Big] = \left(\frac{b}{2} + 2 \sum_{1\le i<j\le b/2} \rho^{j-i}\right)^2 \mathrm{tr}(R_N^2). \tag{S23}$$

An analogous expression holds for the second variance term in Eq. (S21) i.e.

$$\mathrm{Var}\Big[\Big(\sum_{i=b/2+1}^{b} \mathbf{y}_i^c\Big)^\top \Big(\sum_{i=b/2+1}^{b} \hat{\mathbf{y}}_i^c\Big)\Big] = \left(\frac{b}{2} + 2 \sum_{b/2+1\le i<j\le b} \rho^{j-i}\right)^2 \mathrm{tr}(R_N^2) \tag{S24}$$

Let us denote,

$$K = \Big(\sum_{i=1}^{b/2} \mathbf{y}_i^c\Big)^\top \Big(\sum_{i=1}^{b/2} \hat{\mathbf{y}}_i^c\Big), \; \Big(\sum_{i=b/2+1}^{b} \mathbf{y}_i^c\Big)^\top \Big(\sum_{i=b/2+1}^{b} \hat{\mathbf{y}}_i^c\Big)$$

We now expand the covariance term in Eq. (S21). Consider

$$\mathrm{Cov}[K] = \mathbb{E}\Big[\Big(\sum_{i=1}^{b/2} \mathbf{y}_i^c\Big)^\top \Big(\sum_{j=1}^{b/2} \hat{\mathbf{y}}_i^c\Big)\Big(\sum_{k=b/2+1}^{b} \hat{\mathbf{y}}_k^c\Big)^\top \Big(\sum_{m=b/2+1}^{b} \mathbf{y}_m^c\Big)\Big].$$

where $i, j, m$ and $k$ are dummy indices. Using the identity $a^\top b = \mathrm{tr}(ba^\top)$, we rewrite the product as a trace:

$$\begin{aligned}
\mathrm{Cov}[K] &= \mathbb{E}\Big[\,\mathrm{tr}\,\Big(\Big(\sum_{i=1}^{b/2} \mathbf{y}_i^c\Big)\Big(\sum_{m=b/2+1}^{b} \mathbf{y}_m^c\Big)^\top \Big(\sum_{j=1}^{b/2} \hat{\mathbf{y}}_j^c\Big)\Big(\sum_{k=b/2+1}^{b} \hat{\mathbf{y}}_k^c\Big)^\top\Big)\Big] \\
&= \mathrm{tr}\,\Big(\mathbb{E}\Big[\Big(\sum_{i=1}^{b/2}\sum_{m=b/2+1}^{b} \mathbf{y}_i^c \mathbf{y}_m^{c\top}\Big)\Big(\sum_{j=1}^{b/2}\sum_{k=b/2+1}^{b} \hat{\mathbf{y}}_j^c \hat{\mathbf{y}}_k^{c\top}\Big)\Big]\Big) \\
&= \mathrm{tr}\,\Big(\mathbb{E}\Big[\sum_{i=1}^{b/2}\sum_{m=b/2+1}^{b} \mathbf{y}_i^c \mathbf{y}_m^{c\top}\Big]\,\mathbb{E}\Big[\sum_{j=1}^{b/2}\sum_{k=b/2+1}^{b} \hat{\mathbf{y}}_j^c \hat{\mathbf{y}}_k^{c\top}\Big]\Big) \\
&= \mathrm{tr}\,\Big(\sum_{i=1}^{b/2}\sum_{m=b/2+1}^{b} \mathbb{E}[\mathbf{y}_i^c \mathbf{y}_m^{c\top}]\sum_{j=1}^{b/2}\sum_{k=b/2+1}^{b} \mathbb{E}[\hat{\mathbf{y}}_j^c \hat{\mathbf{y}}_k^{c\top}]\Big).
\end{aligned}$$

Using the covariance structure $\mathbb{E}[\mathbf{y}_i^c \mathbf{y}_m^{c\top}] = \mathrm{Cov}[\mathbf{y}_i^c, \mathbf{y}_m^c] = \rho^{m-i} R_N$, we obtain

$$\begin{aligned}
\mathrm{Cov}[K] &= \mathrm{tr}\,\Big(\sum_{i=1}^{b/2}\sum_{m=b/2+1}^{b} \rho^{m-i} R_N \sum_{j=1}^{b/2}\sum_{k=b/2+1}^{b} \rho^{k-j} R_N\Big) \\
&= \mathrm{tr}\,\Big(\Big(\sum_{i=1}^{b/2}\sum_{m=b/2+1}^{b} \rho^{m-i}\Big)\Big(\sum_{j=1}^{b/2}\sum_{k=b/2+1}^{b} \rho^{k-j}\Big) R_N^2\Big) \\
&= \left(\sum_{i=1}^{b/2}\sum_{j=b/2+1}^{b} \rho^{j-i}\right)^2 \mathrm{tr}(R_N^2). \qquad \text{(S25)}
\end{aligned}$$

Substituting Eq. (S23) Eq. (S24) and Eq. (S25) into Eq. (S21), we obtain

$$\mathrm{Var}[P^c_{\mathrm{mes}_1}] = \gamma^2\sigma^4\,\mathrm{tr}(R_N^2)\left[\left(\frac{b}{2} + 2\sum_{1\le i<j\le b/2} \rho^{j-i}\right)^2 + \left(\frac{b}{2} + 2\sum_{b/2+1\le i<j\le b} \rho^{j-i}\right)^2 + 2\left(\sum_{i=1}^{b/2}\sum_{j=b/2+1}^{b} \rho^{j-i}\right)^2\right]$$

By a change of indices, the first two terms are identical, yielding

$$\mathrm{Var}[P^c_{\mathrm{mes}_1}] = 2\gamma^2\sigma^4\,\mathrm{tr}(R_N^2).\left[\left(\frac{b}{2} + 2\sum_{1\le i<j\le b/2} \rho^{j-i}\right)^2 + \left(\sum_{i=1}^{b/2}\sum_{j=b/2+1}^{b} \rho^{j-i}\right)^2\right]. \tag{S26}$$

Following the same derivation steps as in the microscopic case (see Eq. (S5)), the first summation term in Eq. (S26) is obtained.

$$\sum_{1\le i<j\le b/2} \rho^{j-i} = \frac{(b/2-1)\rho - (b/2)\rho^2 + \rho^{b/2+1}}{(1-\rho)^2} = G. \tag{S27}$$

We now simplify the second sum in Eq. (S26), by separating the dependence on $i$ and $j$:

$$\sum_{i=1}^{b/2}\sum_{j=b/2+1}^{b} \rho^{j-i} = \sum_{i=1}^{b/2}\rho^{-i}\sum_{j=b/2+1}^{b}\rho^{j}. \tag{S28}$$

Next, we evaluate the inner sum. Using a change of variable $j = j' + b/2$, where $j' = 1, \ldots, b/2$, we obtain

$$\sum_{j=b/2+1}^{b}\rho^{j} = \sum_{j'=1}^{b/2}\rho^{j'+b/2} = \rho^{b/2}\sum_{j'=1}^{b/2}\rho^{j'}.$$

Substituting above expression in Eq. (S28),

$$\sum_{i=1}^{b/2}\sum_{j=b/2+1}^{b} \rho^{j-i} = \rho^{b/2}\left(\sum_{i=1}^{b/2}\rho^{-i}\right)\left(\sum_{j'=1}^{b/2}\rho^{j'}\right). \tag{S29}$$

We now evaluate both sums using the finite geometric series formula. First,

$$\sum_{j'=1}^{b/2}\rho^{j'} = \frac{\rho(1-\rho^{b/2})}{1-\rho}.$$

Similarly,

$$\sum_{i=1}^{b/2}\rho^{-i} = \frac{\rho^{-1}(1-\rho^{-b/2})}{1-\rho^{-1}}.$$

Substituting these expressions in Eq. (S29), we obtain

$$\sum_{i=1}^{b/2}\sum_{j=b/2+1}^{b} \rho^{j-i} = \rho^{b/2}\left(\frac{\rho^{-1}(1-\rho^{-b/2})}{1-\rho^{-1}}\right)\left(\frac{\rho(1-\rho^{b/2})}{1-\rho}\right).$$

Which upon simplifying, yields,

$$\sum_{i=1}^{b/2}\sum_{j=b/2+1}^{b}\rho^{j-i}=\frac{\rho(1-\rho^{b/2})^2}{(1-\rho)^2}. \tag{S30}$$

Substituting values of Eq. (S27) and Eq. (S30) into Eq. (S26), we obtain,

$$\mathrm{Var}[P^c_{\mathrm{mes}_1}]=2\sigma^4\gamma^2\,\mathrm{tr}(R_N^2)\left[\left(\frac{b}{2}+2G\right)^2+\frac{\rho^2(1-\rho^{b/2})^4}{(1-\rho)^4}\right]. \tag{S31}$$

By identical arguments, $\mathrm{Var}[P^{\mathrm{x}}_{\mathrm{mes}_1}]$ has the same form.

$$\mathrm{Var}[P^{\mathrm{x}}_{\mathrm{mes}_1}]=2\sigma^4\gamma^2\,\mathrm{tr}(R_N^2)\left[\left(\frac{b}{2}+2G\right)^2+\frac{\rho^2(1-\rho^{b/2})^4}{(1-\rho)^4}\right]. \tag{S32}$$

Substituting expectations Eq. (S19) Eq. (S20) and variances Eq. (S31) Eq. (S32) into the definition of the $d'$ ( Eq. (4) in main text) yields

$$d'_{\mathrm{mes}_1}=\frac{\sum_{i=1}^{b/2}\sum_{j=1}^{b/2}\left(\langle\boldsymbol{\mu}_i^c,\boldsymbol{\mu}_j^c\rangle-\langle\boldsymbol{\mu}_i^c,\boldsymbol{\mu}_j^{c'}\rangle\right)+\sum_{i=b/2+1}^{b}\sum_{j=b/2+1}^{b}\left(\langle\boldsymbol{\mu}_i^c,\boldsymbol{\mu}_j^c\rangle-\langle\boldsymbol{\mu}_i^c,\boldsymbol{\mu}_j^{c'}\rangle\right)}{\sqrt{2\sigma^4\,\mathrm{tr}(R_N^2)\left[\left(\frac{b}{2}+2G\right)^2+\rho^2\left(\frac{1-\rho^{b/2}}{1-\rho}\right)^4\right]}}. \qquad\square$$

#### Sensitivity Index at the Macroscale

PROOF OF THEOREM 4. Let $\mathbf{v}^c_{\mathrm{mac}}$ denote the temporally encoded trial vector at macroscale resolution:

$$\mathbf{v}^c_{\mathrm{mac}}=\mathbf{S}\Big(\sum_{i=1}^{b}(\boldsymbol{\mu}_i^c+\sigma\boldsymbol{y}_i^{j,c})\Big)\odot\mathbf{t}. \tag{S33}$$

Let $T=\mathrm{diag}(\mathbf{t})$. Since $T^\top T=I$ and $\mathbf{S}^\top\mathbf{S}\approx\gamma I$, the within-class inner product simplifies to

$$P^c_{\mathrm{mac}}=\gamma\Big(\sum_{i=1}^{b}(\boldsymbol{\mu}_i^c+\sigma\mathbf{y}_i^c)\Big)^\top\Big(\sum_{k=1}^{b}(\boldsymbol{\mu}_k^c+\sigma\hat{\mathbf{y}}_k^c)\Big).$$

Using $\mathbb{E}[\mathbf{y}_i^c]=0$, we obtain within class expectation as,

$$\mathbb{E}[P^c_{\mathrm{mac}}]=\gamma\sum_{i=1}^{b}\sum_{k=1}^{b}\langle\boldsymbol{\mu}_i^c,\boldsymbol{\mu}_k^c\rangle. \tag{S34}$$

Similarly, for between-class,

$$\mathbb{E}[P^{\mathrm{x}}_{\mathrm{mac}}]=\gamma\sum_{i=1}^{b}\sum_{k=1}^{b}\langle\boldsymbol{\mu}_i^c,\boldsymbol{\mu}_k^{c'}\rangle. \tag{S35}$$

We now compute the variance of the within-class similarity:

$$\mathrm{Var}[P^c_{\mathrm{mac}}] = \gamma^2 \mathrm{Var}\Big[\sum_{i=1}^{b}\sum_{k=1}^{b}(\boldsymbol{\mu}^c_i + \sigma \mathbf{y}^c_i)^\top(\boldsymbol{\mu}^c_k + \sigma \hat{\mathbf{y}}^c_k)\Big].$$

Using statistical model properties, only the quadratic noise term contributes:

$$\mathrm{Var}[P^c_{\mathrm{mac}}] = \gamma^2\sigma^4 \mathrm{Var}\Big[\Big(\sum_{i=1}^{b}\mathbf{y}^c_i\Big)^\top\Big(\sum_{k=1}^{b}\hat{\mathbf{y}}^c_k\Big)\Big].$$

Using the trace formulation,

$$\mathrm{Var}\Big[\Big(\sum_{i=1}^{b}\mathbf{y}^c_i\Big)^\top\Big(\sum_{i=1}^{b}\hat{\mathbf{y}}^c_i\Big)\Big] = \mathrm{tr}\left(\mathrm{Cov}\left[\sum_{i=1}^{b}\mathbf{y}^c_i\right]\mathrm{Cov}\left[\sum_{i=1}^{b}\hat{\mathbf{y}}^c_i\right]\right). \tag{S36}$$

We now compute the covariance in Eq. (S36):

$$\begin{aligned}\mathrm{Cov}\left[\sum_{i=1}^{b}\mathbf{y}^c_i\right] &= \mathbb{E}\Big[\Big(\sum_{i=1}^{b}\mathbf{y}^c_i\Big)\Big(\sum_{j=1}^{b}\mathbf{y}^c_j\Big)^\top\Big] \\ &= \sum_{i=1}^{b}\sum_{j=1}^{b}\mathbb{E}[\mathbf{y}^c_i\mathbf{y}^{c\top}_j] \\ &= \sum_{i=1}^{b}\sum_{j=1}^{b}\mathrm{Cov}[\mathbf{y}^c_i,\mathbf{y}^c_j] \\ &= bR_N + 2\sum_{1\le i<j\le b}\rho^{j-i}R_N.\end{aligned}$$

An identical expression holds for the $\mathrm{Cov}\left[\sum_{i=1}^{b}\hat{\mathbf{y}}^c_i\right]$. Substituting these covariance expansions back into Eq. (S36), we obtain

$$\mathrm{Var}\Big[\Big(\sum_{i=1}^{b}\mathbf{y}^c_i\Big)^\top\Big(\sum_{i=1}^{b}\hat{\mathbf{y}}^c_i\Big)\Big] = \mathrm{tr}(R_N^2)\left(b + 2\sum_{1\le i<j\le b}\rho^{j-i}\right)^2.$$

Following the similar derivation steps in Eq. (S5) the geometric sum $\sum_{1\le i<j\le b}\rho^{j-i}$ can be defined as

$$\sum_{1\le i<j\le b}\rho^{j-i} = \frac{(b-1)\rho - b\rho^2 + \rho^{b+1}}{(1-\rho)^2} = F.$$

Thus,

$$\mathrm{Var}[P^c_{\mathrm{mac}}] = \sigma^4\gamma^2\,\mathrm{tr}(R_N^2)(b+2F)^2. \tag{S37}$$

By identical arguments,

$$\mathrm{Var}[P^{\mathrm{x}}_{\mathrm{mac}}] = \sigma^4\gamma^2\,\mathrm{tr}(R_N^2)(b+2F)^2. \tag{S38}$$

Substituting expectations in Eq. (S34) Eq. (S35) and variances in Eq. (S37) Eq. (S38) into the definition of $d'$ ( Eq. (4) in main text) yields,

$$d'_{\mathrm{mac}} = \frac{\sum_{i=1}^{b}\sum_{j=1}^{b}\Big(\langle\boldsymbol{\mu}_i^c,\boldsymbol{\mu}_j^c\rangle - \langle\boldsymbol{\mu}_i^c,\boldsymbol{\mu}_j^{c'}\rangle\Big)}{\sqrt{\sigma^4\,\mathrm{tr}(R_N^2)(b+2F)^2}}. \qquad \square$$

### SUPPLEMENTARY NOTE 2: EXAMPLE GENERALIZATION TO CORRELATED SIGNAL AND NOISE

In this Supplementary Note we provide an example for how our results in the main text can be generalized if Assumption 1 does not hold. Because relaxing this assumption significantly increases the complexity of deriving closed-form expressions, we only analyze a *minimal setting* consisting of two time points ($b = 2$), with scaling factor ($k = 1$) and independence among neurons ($R_N = I_N$), but other aspects of the statistical model remain general.

For clarity and lack of confusion with the triad of scales in the main text, we refer to the microscale (no temporal averaging) and macroscale (full temporal averaging) regimes as the *before-averaging* and *after-averaging* regimes, respectively. To better understand the effects of Assumption 1, we compare the resulting decodability expressions both with and without this assumption. Under Assumption 1 (simplified model), it immediately follows from Theorems 1 and 4 in the main text that the corresponding sensitivity indices take the form,

$$d'_{\mathrm{bef}} = \frac{\|\boldsymbol{\mu}_1^c\|^2 + \|\boldsymbol{\mu}_2^c\|^2 - \langle\boldsymbol{\mu}_1^c,\boldsymbol{\mu}_1^{c'}\rangle - \langle\boldsymbol{\mu}_2^c,\boldsymbol{\mu}_2^{c'}\rangle}{\sqrt{2\sigma^4 N(1+\rho^2)}}, \tag{S39}$$

$$d'_{\mathrm{aft}} = \frac{\|\boldsymbol{\mu}_1^c\|^2 + \|\boldsymbol{\mu}_2^c\|^2 + 2\langle\boldsymbol{\mu}_1^c,\boldsymbol{\mu}_2^c\rangle - \langle\boldsymbol{\mu}_1^c,\boldsymbol{\mu}_1^{c'}\rangle - \langle\boldsymbol{\mu}_2^c,\boldsymbol{\mu}_2^{c'}\rangle - \langle\boldsymbol{\mu}_1^c,\boldsymbol{\mu}_2^{c'}\rangle - \langle\boldsymbol{\mu}_2^c,\boldsymbol{\mu}_1^{c'}\rangle}{\sqrt{4\sigma^4 N(1+\rho)^2}}. \tag{S40}$$

#### *Before-Averaging Sensitivity Index in the Presence of Signal–Noise Correlations*

We now relax Assumption 1 and derive the corresponding sensitivity index expression for the before-averaging regime under the general two-time points setting.

THEOREM S1. *Let*

$$\mathbf{v}^c_{\mathrm{bef}} = \mathbf{S}\mathbf{n}^c_1 \odot \mathbf{t}_1 + \mathbf{S}\mathbf{n}^c_2 \odot \mathbf{t}_2,$$

*where* $\mathbf{n}^c_1$ *and* $\mathbf{n}^c_2$ *follows the statistical model in Eq. (5), and let us denote the within-class and between-class similarities, as*

$$P^c_{\mathrm{bef}} = {\mathbf{v}^c_{\mathrm{bef}}}^\top \hat{\mathbf{v}}^c_{\mathrm{bef}}, \qquad P^{\mathrm{x}}_{\mathrm{bef}} = {\mathbf{v}^c_{\mathrm{bef}}}^\top \hat{\mathbf{v}}^{c'}_{\mathrm{bef}}, \quad \textit{for} \quad c' \neq c, \tag{S41}$$

*Then*

$$d'_{\mathrm{bef}} = \frac{S_{\mathrm{bef}}}{\sqrt{N_{\mathrm{bef}}}},$$

*where*

$$S_{\text{bef}} = \|\boldsymbol{\mu}_1^c\|^2 + \|\boldsymbol{\mu}_2^c\|^2 - \langle \boldsymbol{\mu}_1^c, \boldsymbol{\mu}_1^{c'} \rangle - \langle \boldsymbol{\mu}_2^c, \boldsymbol{\mu}_2^{c'} \rangle,$$

*and*

$$N_{\text{bef}} = \sigma^2 \Big[ \frac{3}{2}\|\boldsymbol{\mu}_1^{c'}\|^2 + \frac{3}{2}\|\boldsymbol{\mu}_1^c\|^2 + \frac{1}{2}\|\boldsymbol{\mu}_2^{c'}\|^2 + \frac{1}{2}\|\boldsymbol{\mu}_2^c\|^2$$
$$+ \rho \left( \langle \boldsymbol{\mu}_1^{c'}, \boldsymbol{\mu}_2^{c'} \rangle + 3\langle \boldsymbol{\mu}_1^c, \boldsymbol{\mu}_2^c \rangle \right) + 2\sigma^2 N(1+\rho^2) \Big].$$

PROOF. Let $T_1 = \text{diag}(\mathbf{t}_1)$ and $T_2 = \text{diag}(\mathbf{t}_2)$. Since the temporal basis vectors are orthogonal, we have $T_1^\top T_2 = 0$ and $T_i^\top T_i = I$. Hence, within class inner product simplifies as

$$P_{\text{bef}}^c = \gamma(\mathbf{n}_1^\top \hat{\mathbf{n}}_1 + \mathbf{n}_2^\top \hat{\mathbf{n}}_2). \tag{S42}$$

Since neural responses follows the statistical model in Eq. (5) we obtain,

$$\begin{aligned} P_{\text{bef}}^c &= \gamma((\boldsymbol{\mu}_1^c + \sigma \mathbf{y}_1^c)^\top (\boldsymbol{\mu}_1^c + \sigma \hat{\mathbf{y}}_1^c) + (\boldsymbol{\mu}_2^c + \sigma \mathbf{y}_2^c)^\top (\boldsymbol{\mu}_2^c + \sigma \hat{\mathbf{y}}_2^c)) \\ &= \gamma(\|\boldsymbol{\mu}_1^c\|^2 + \sigma \mathbf{y}_1^{c\top} \boldsymbol{\mu}_1^c + \sigma \boldsymbol{\mu}_1^{c\top} \hat{\mathbf{y}}_1^c + \sigma^2 \mathbf{y}_1^{c\top} \hat{\mathbf{y}}_1^c \\ &\quad + \|\boldsymbol{\mu}_2^c\|^2 + \sigma \mathbf{y}_2^{c\top} \boldsymbol{\mu}_2^c + \sigma \boldsymbol{\mu}_2^{c\top} \hat{\mathbf{y}}_2^c + \sigma^2 \mathbf{y}_2^{c\top} \hat{\mathbf{y}}_2^c). \end{aligned}$$

Hence the expectation of within class similarities would be,

$$\mathbb{E}[P_{\text{bef}}^c] = \gamma(\|\boldsymbol{\mu}_1^c\|^2 + \|\boldsymbol{\mu}_2^c\|^2). \tag{S43}$$

Similarly, for between class similarities

$$\mathbb{E}[P_{\text{bef}}^{\text{x}}] = \gamma(\langle \boldsymbol{\mu}_1^c, \boldsymbol{\mu}_1^{c'} \rangle + \langle \boldsymbol{\mu}_2^c, \boldsymbol{\mu}_2^{c'} \rangle). \tag{S44}$$

We now compute variance of within-class similarity. Ignoring constant signal terms, we write

$$\text{Var}[P_{\text{bef}}^c] = \gamma^2(\sigma^2 \text{Var}[A] + \sigma^4 \text{Var}[B]), \tag{S45}$$

where

$$\begin{aligned} A &= \mathbf{y}_1^{c\top} \boldsymbol{\mu}_1^c + \boldsymbol{\mu}_1^{c\top} \hat{\mathbf{y}}_1^c + \mathbf{y}_2^{c\top} \boldsymbol{\mu}_2^c + \boldsymbol{\mu}_2^{c\top} \hat{\mathbf{y}}_2^c, \\ B &= \mathbf{y}_1^{c\top} \hat{\mathbf{y}}_1^c + \mathbf{y}_2^{c\top} \hat{\mathbf{y}}_2^c. \end{aligned}$$

We evaluate each variance term in Eq. (S45) separately. First we expand quadratic noise term $\text{Var}[B]$,

$$\text{Var}[B] = \mathbb{E}\big[\mathbf{y}_1^{c\top} \hat{\mathbf{y}}_1^c \mathbf{y}_1^{c\top} \hat{\mathbf{y}}_1^c + \mathbf{y}_2^{c\top} \hat{\mathbf{y}}_2^c \mathbf{y}_2^{c\top} \hat{\mathbf{y}}_2^c + 2\mathbf{y}_1^{c\top} \hat{\mathbf{y}}_1^c \mathbf{y}_2^{c\top} \hat{\mathbf{y}}_2^c\big].$$

Using trace representation and independence across trials,

$$\text{Var}[B] = 2N(1+\rho^2). \tag{S46}$$

Now expanding signal-noise term $\text{Var}[A]$,

$$\text{Var}[A] = \sum \text{Var(individual terms)} + 2\sum \text{Cov(cross terms)}. \tag{S47}$$

Each variance term in Eq. (S47) simplifies as

$$\mathrm{Var}[\boldsymbol{\mu}_1^{c\top}\hat{\mathbf{y}}_1^c] = \mathbb{E}[\boldsymbol{\mu}_1^{c\top}\hat{\mathbf{y}}_1^c\hat{\mathbf{y}}_1^{c\top}\boldsymbol{\mu}_1^c] \tag{S48a}$$

$$= \boldsymbol{\mu}_1^{c\top} I_N \boldsymbol{\mu}_1^c \tag{S48b}$$

$$= \|\boldsymbol{\mu}_1^c\|^2 = \mathrm{Var}[\mathbf{y}_1^{c\top}\boldsymbol{\mu}_1^c] \tag{S48c}$$

Similar to derivation in Eq. (S48) we have,

$$\mathrm{Var}[\mathbf{y}_2^{c\top}\boldsymbol{\mu}_2^c] = \mathrm{Var}[\boldsymbol{\mu}_2^{c\top}\hat{\mathbf{y}}_2^c] = \|\boldsymbol{\mu}_2^c\|^2 \tag{S49}$$

The covariance terms in Eq. (S47) expanded using $\mathrm{Cov}[a,b] = \mathbb{E}[ab^\top]$ for $a, b$ be a mean zero random vectors, and similar derivation steps in Eq. (S48), evaluate to

$$\mathrm{Cov}[\boldsymbol{\mu}_1^{c\top}\hat{\mathbf{y}}_1^c, \boldsymbol{\mu}_2^{c\top}\hat{\mathbf{y}}_2^c] = \rho\langle\boldsymbol{\mu}_1^c, \boldsymbol{\mu}_2^c\rangle = \mathrm{Cov}[\mathbf{y}_1^{c\top}\boldsymbol{\mu}_1^c, \mathbf{y}_2^{c\top}\boldsymbol{\mu}_2^c] \tag{S50}$$

Collecting all contributions of variances Eq. (S48), Eq. (S49) and covariances Eq. (S50) and substituting back in Eq. (S47),

$$\mathrm{Var}[A] = 2\left(\|\boldsymbol{\mu}_1^c\|^2 + \|\boldsymbol{\mu}_2^c\|^2 + 2\rho\langle\boldsymbol{\mu}_1^c, \boldsymbol{\mu}_2^c\rangle\right). \tag{S51}$$

Substituting Eq. (S51) and Eq. (S46) into Eq. (S45), we obtain

$$\mathrm{Var}[P_{\text{bef}}^c] = 2\gamma^2(\sigma^2\left[\|\boldsymbol{\mu}_1^c\|^2 + \|\boldsymbol{\mu}_2^c\|^2 + 2\rho\langle\boldsymbol{\mu}_1^c, \boldsymbol{\mu}_2^c\rangle\right] + 2\sigma^4 N(1+\rho^2)). \tag{S52}$$

To compute between class similarities variance, proceeding analogously, and following the same expansions, we obtain

$$\mathrm{Var}[P_{\text{bef}}^{\text{x}}] = \sigma^2\gamma^2\Big[\|\boldsymbol{\mu}_1^{c'}\|^2 + \|\boldsymbol{\mu}_1^c\|^2 + \|\boldsymbol{\mu}_2^{c'}\|^2 + \|\boldsymbol{\mu}_2^c\|^2 + 2\rho\left(\langle\boldsymbol{\mu}_1^{c'}, \boldsymbol{\mu}_2^{c'}\rangle + \langle\boldsymbol{\mu}_1^c, \boldsymbol{\mu}_2^c\rangle\right)$$
$$+2\sigma^4 N(1+\rho^2)\Big]. \tag{S53}$$

Substituting the expectations ( Eq. (S43) and Eq. (S44)) and variances ( Eq. (S52) and Eq. (S53)) into the definition of $d'$( Eq. (4) in main text) yields the desired expression. □

### *After-Averaging Sensitivity Index in the Presence of Signal–Noise Correlations*

We now derive the corresponding sensitivity index expression for the after-averaging regime in the minimal (two-bin) setting without enforcing Assumption 1.

THEOREM S2. *Let*

$$\mathbf{v}_{\text{aft}}^c = (\mathbf{n}_1^c + \mathbf{n}_2^c)\mathbf{t}$$

*where $\mathbf{n}_1^c$ and $\mathbf{n}_2^c$ follows statistical model in Eq. (5), and let us denote the within-class and between-class similarities, as*

$$P_{\text{aft}}^c = \mathbf{v}_{\text{aft}}^{c\;\top}\hat{\mathbf{v}}_{\text{aft}}^c, \qquad P_{\text{aft}}^{\text{x}} = \mathbf{v}_{\text{aft}}^{c\;\top}\hat{\mathbf{v}}_{\text{aft}}^{c'}, \quad \textit{for} \quad c' \neq c, \tag{S54}$$

*Then*

$$d'_{\text{aft}} = \frac{N_{\text{aft}}}{\sqrt{D_{\text{aft}}}},$$

*where*

$$N_{\text{aft}} = \|\boldsymbol{\mu}_1^c\|^2 + \|\boldsymbol{\mu}_2^c\|^2 + 2\langle \boldsymbol{\mu}_1^c, \boldsymbol{\mu}_2^c \rangle - \langle \boldsymbol{\mu}_1^c, \boldsymbol{\mu}_1^{c'} \rangle - \langle \boldsymbol{\mu}_2^c, \boldsymbol{\mu}_2^{c'} \rangle - \langle \boldsymbol{\mu}_1^c, \boldsymbol{\mu}_2^{c'} \rangle - \langle \boldsymbol{\mu}_2^c, \boldsymbol{\mu}_1^{c'} \rangle,$$

*and*

$$D_{\text{aft}} = \sigma^2 \Big[ (1+\rho)\big(\|\boldsymbol{\mu}_1^{c'}\|^2 + \|\boldsymbol{\mu}_2^{c'}\|^2\big) + (3+\rho)\big(\|\boldsymbol{\mu}_1^c\|^2 + \|\boldsymbol{\mu}_2^c\|^2\big)$$
$$+ 2(2+3\rho)\langle \boldsymbol{\mu}_1^c, \boldsymbol{\mu}_2^c \rangle + \rho \langle \boldsymbol{\mu}_1^{c'}, \boldsymbol{\mu}_2^{c'} \rangle + 4\sigma^2 N (1+\rho)^2 \Big].$$

PROOF. The after-averaging trial vectors within class similiraty can be written as

$$P_{\text{aft}}^c = \gamma(\boldsymbol{\mu}_1^c + \sigma \mathbf{y}_1^c + \boldsymbol{\mu}_2^c + \sigma \mathbf{y}_2^c)^\top (\boldsymbol{\mu}_1^c + \sigma \hat{\mathbf{y}}_1^c + \boldsymbol{\mu}_2^c + \sigma \hat{\mathbf{y}}_2^c).$$

Expanding and taking expectations using $\mathbb{E}[\mathbf{y}_i^c] = 0$, we obtain the within-class expectation

$$\mathbb{E}[P_{\text{aft}}^c] = \gamma(\|\boldsymbol{\mu}_1^c\|^2 + \|\boldsymbol{\mu}_2^c\|^2 + 2\langle \boldsymbol{\mu}_1^c, \boldsymbol{\mu}_2^c \rangle), \tag{S55}$$

and the between-class expectation

$$\mathbb{E}[P_{\text{aft}}^{\text{x}}] = \gamma(\langle \boldsymbol{\mu}_1^c, \boldsymbol{\mu}_1^{c'} \rangle + \langle \boldsymbol{\mu}_2^c, \boldsymbol{\mu}_2^{c'} \rangle + \langle \boldsymbol{\mu}_1^c, \boldsymbol{\mu}_2^{c'} \rangle + \langle \boldsymbol{\mu}_2^c, \boldsymbol{\mu}_1^{c'} \rangle). \tag{S56}$$

We now compute the variance of the within-class similarity. Grouping signal–noise and noise–noise contributions yields

$$\begin{aligned}
\text{Var}[P_{\text{aft}}^c] = \sigma^2\gamma^2 \Big( & \text{Var}\Big[\boldsymbol{\mu}_1^{c\top}(\hat{\mathbf{y}}_1^c + \hat{\mathbf{y}}_2^c)\Big] + \text{Var}\Big[\boldsymbol{\mu}_2^{c\top}(\hat{\mathbf{y}}_1^c + \hat{\mathbf{y}}_2^c)\Big] \\
& + 2\ \text{Cov}\Big[\boldsymbol{\mu}_1^{c\top}(\hat{\mathbf{y}}_1^c + \hat{\mathbf{y}}_2^c)\,, \boldsymbol{\mu}_2^{c\top}(\hat{\mathbf{y}}_1^c + \hat{\mathbf{y}}_2^c)\Big] \\
& + \text{Var}\Big[\mathbf{y}_1^{c\top}(\boldsymbol{\mu}_1^c + \boldsymbol{\mu}_2^c)\Big] + \text{Var}\Big[\mathbf{y}_2^{c\top}(\boldsymbol{\mu}_1^c + \boldsymbol{\mu}_2^c)\Big] \\
& + 2\ \text{Cov}\Big[\mathbf{y}_1^{c\top}(\boldsymbol{\mu}_1^c + \boldsymbol{\mu}_2^c)\,, \mathbf{y}_2^{c\top}(\boldsymbol{\mu}_1^c + \boldsymbol{\mu}_2^c)\Big] \\
& + \text{Var}\Big[(\mathbf{y}_1^c + \mathbf{y}_2^c)^\top(\hat{\mathbf{y}}_1^c + \hat{\mathbf{y}}_2^c)\Big] \Big).
\end{aligned} \tag{S57}$$

The individual variance terms of signal-noise contributions in Eq. (S57) simplify using similar derivation steps used in Eq. (S48). In particular, we have the following simplifications,

$$\text{Var}\Big[\boldsymbol{\mu}_1^{c\top}(\hat{\mathbf{y}}_1^c + \hat{\mathbf{y}}_2^c)\Big] = 2\|\boldsymbol{\mu}_1^c\|^2, \qquad \text{Var}\Big[\boldsymbol{\mu}_2^{c\top}(\hat{\mathbf{y}}_1^c + \hat{\mathbf{y}}_2^c)\Big] = 2\|\boldsymbol{\mu}_2^c\|^2 \tag{S58a}$$

$$\text{Var}\Big[\mathbf{y}_1^{c\top}(\boldsymbol{\mu}_1^c + \boldsymbol{\mu}_2^c)\Big] = \text{Var}\Big[\mathbf{y}_2^{c\top}(\boldsymbol{\mu}_1^c + \boldsymbol{\mu}_2^c)\Big] = \|\boldsymbol{\mu}_1^c\|^2 + \|\boldsymbol{\mu}_2^c\|^2 \tag{S58b}$$

The signal driven covariance term in Eq. (S57) can be simplified as

$$\text{Cov}\Big[\boldsymbol{\mu}_1^{c\top}(\hat{\mathbf{y}}_1^c + \hat{\mathbf{y}}_2^c)\,, \boldsymbol{\mu}_2^{c\top}(\hat{\mathbf{y}}_1^c + \hat{\mathbf{y}}_2^c)\Big] = \mathbb{E}\Big[\boldsymbol{\mu}_1^{c\top}(\hat{\mathbf{y}}_1^c + \hat{\mathbf{y}}_2^c)(\hat{\mathbf{y}}_1^c + \hat{\mathbf{y}}_2^c)^\top \boldsymbol{\mu}_2^c\Big]$$

$$= \boldsymbol{\mu}_1^{c\top}\mathbb{E}[I_N + I_N + \rho I_N + \rho I_N]\boldsymbol{\mu}_2^c$$
$$= 2(1+\rho)\langle \boldsymbol{\mu}_1^c, \boldsymbol{\mu}_2^c \rangle, \tag{S59}$$

similarly for the noise-driven covariace term in Eq. (S57) we have

$$\mathrm{Cov}\Big[\mathbf{y}_1^{c\top}(\boldsymbol{\mu}_1^c + \boldsymbol{\mu}_2^c)\,, \mathbf{y}_2^{c\top}(\boldsymbol{\mu}_1^c + \boldsymbol{\mu}_2^c)\Big] = \rho(\|\boldsymbol{\mu}_1^c\|^2 + \|\boldsymbol{\mu}_2^c\|^2 + \langle \boldsymbol{\mu}_1^c, \boldsymbol{\mu}_2^c \rangle). \tag{S60}$$

The variance term with pure noise contribution in Eq. (S57) evaluates to

$$\mathrm{Var}\left[(\mathbf{y}_1^c + \mathbf{y}_2^c)^\top (\hat{\mathbf{y}}_1^c + \hat{\mathbf{y}}_2^c)\right] = 4\sigma^4 N(1+\rho)^2. \tag{S61}$$

Substituting these components(i.e. Eq. (S58) Eq. (S59) Eq. (S60) and Eq. (S61)) into Eq. (S57) yields the following expression for the within-class variance,

$$\mathrm{Var}[P_{\text{bef}}^c] = \sigma^2\gamma^2\left[4\|\boldsymbol{\mu}_1^c\|^2 + 4\|\boldsymbol{\mu}_2^c\|^2 + 2\Big(2(1+2\rho)\langle \boldsymbol{\mu}_1^c, \boldsymbol{\mu}_2^c \rangle + \rho(\|\boldsymbol{\mu}_1^c\|^2 + \|\boldsymbol{\mu}_2^c\|^2)\Big) + 4\sigma^2 N(1+\rho)^2\right]. \tag{S62}$$

Now, for the between-class similarity, proceeding under the same decomposition for $P_{\text{aft}}^{\text{x}}$, as in Eq. (S57), and by simplifying each term analogously, we obtain

$$\mathrm{Var}[P_{\text{aft}}^{\text{x}}] = \sigma^2\gamma^2\Big[\|\boldsymbol{\mu}_1^{c'}\|^2 + \|\boldsymbol{\mu}_1^c\|^2 + \|\boldsymbol{\mu}_2^{c'}\|^2 + \|\boldsymbol{\mu}_2^c\|^2 + 2(1+\rho)\langle \boldsymbol{\mu}_1^c, \boldsymbol{\mu}_2^c \rangle$$
$$+\rho(\|\boldsymbol{\mu}_1^{c'}\|^2 + \|\boldsymbol{\mu}_2^{c'}\|^2 + 2\langle \boldsymbol{\mu}_1^{c'}, \boldsymbol{\mu}_2^{c'} \rangle) + 2\sigma^2 N(1+\rho)^2\Big]. \tag{S63}$$

Thus, both within-class and between-class variances are fully characterized.

Substituting the corresponding expectations( Eq. (S55), Eq. (S56)) and variances( Eq. (S62), Eq. (S63)) into the definition of the sensitivity index $d'$( Eq. (4) in main text) yields the desired expression. □

#### *Determining the Optimal Resolution in the Absence of Signal–Noise Correlations*

We now compare the sensitivity indices before and after averaging in the above two-bin setting when the signal–noise correlations are assumed to be zero (i.e., when Assumption 1 holds). Using the same inner-product structure on signal representations as in Eq. (7b), with $\|\boldsymbol{\mu}_i^c\| = \nu$ for $i = 1, 2$, the sensitivity indices in Eq. (S39) and Eq. (S40) reduces to

$$d'_{\text{bef}} = \frac{2\nu^2}{\sigma^2\sqrt{2(1+\rho^2)N}}, \qquad d'_{\text{aft}} = \frac{2\nu^2(1+\beta)}{\sigma^2\sqrt{4(1+\rho)^2\,N}}.$$

To determine when temporal averaging improves discriminability, we compare the two regimes via the inequality $d'_{\text{aft}} > d'_{\text{bef}}$. This yields

$$\frac{2\nu^2(1+\beta)}{\sigma^2\sqrt{4(1+\rho)^2\,N}} > \frac{2\nu^2}{\sigma^2\sqrt{2(1+\rho^2)N}}$$
$$\Leftrightarrow (1+\beta)^2 > \frac{2(1+\rho)^2}{1+\rho^2}. \tag{S64}$$

The corresponding optimality function is therefore,

$$\mathcal{G}_2^{\mathrm{aft>bef}} = (1+\beta)^2 - \frac{2(1+\rho)^2}{1+\rho^2}. \tag{S65}$$

Temporal averaging is beneficial in this regime whenever $\mathcal{G}_2^{\mathrm{aft>bef}} > 0$. The corresponding before- and after-averaging regimes are illustrated in the "simplified model" column of Figure Eq. (S1), which summarizes the main qualitative differences induced by temporal averaging.

#### *Determining the Optimal Resolution in the Presence of Signal–Noise Correlations.*

We now extend the comparison to the general setting where signal–noise correlations are non-zero. Using the derived sensitivity index expressions for the two-bin case in Theorem S1 and Thoerem S2 with the same inner-product structure in Eq. (7b), we obtain

$$\begin{aligned} d'_{\mathrm{bef}} &= \frac{2\nu^2}{\sqrt{N\sigma^2\left(4\nu^2 + 4\rho\nu^2\beta + 2\sigma^2(1+\rho^2)\right)}}, \\ d'_{\mathrm{aft}} &= \frac{2\nu^2(1+\beta)}{\sqrt{N\sigma^2\left(8\nu^2 + (4\nu^2 + 8\rho\nu^2)\,\beta + 4\nu^2\rho + 4\sigma^2(1+\rho)^2\right)}}. \end{aligned}$$

which, in comparison, gives

$$\begin{aligned} & d'_{\mathrm{aft}} > d'_{\mathrm{bef}} \Leftrightarrow \\ & \frac{2\nu^2(1+\beta)}{\sqrt{N\sigma^2\left(8\nu^2 + (4\nu^2 + 8\rho\nu^2)\,\beta + 4\nu^2\rho + 4\sigma^2(1+\rho)^2\right)}} > \frac{2\nu^2}{\sqrt{N\sigma^2\left(4\nu^2 + 4\rho\nu^2\beta + 2\sigma^2(1+\rho^2)\right)}}. \end{aligned}$$

For notational convenience, we define the optimality function

$$\mathcal{H}_2^{\mathrm{aft>bef}} = \frac{(1+\beta)^2}{8\nu^2 + (4\nu^2 + 8\rho\nu^2)\,\beta + 4\nu^2\rho + 4\sigma^2(1+\rho)^2} - \frac{1}{4\nu^2 + 4\rho\nu^2\beta + 2\sigma^2(1+\rho^2)}. \tag{S66}$$

Thus, temporal averaging improves discriminability in the general setting whenever $\mathcal{H}_2^{\mathrm{aft>bef}} > 0$.

We compare both the simplified and the general formulations by visualizing the corresponding optimality functions in Figure S1. These comparisons highlight how the relative magnitudes of the signal parameter $\nu$ and noise parameter $\sigma$ shape the optimal temporal scale. In particular, since $\nu$ and $\sigma$ explicitly influence only the general model, the two formulations coincide in the high-noise regime ($\nu \ll \sigma$)( Fig. S1(b)). In this case, both $\mathcal{G}_2^{\mathrm{aft>bef}}$ and $\mathcal{H}_2^{\mathrm{aft>bef}}$ yield consistent boundaries (i.e. $\mathcal{G}_2^{\mathrm{aft>bef}} = \mathcal{H}_2^{\mathrm{aft>bef}} = 0$ ), confirming that the simplified analysis accurately captures the behavior of the full model when magnitude of noise dominates. In contrast, in the high-signal regime ($\nu \gg \sigma$) ( Fig. S1(c)), averaging becomes increasingly favorable, leading to a broader region where the after-averaging (macroscale) representation is optimal. For intermediate regimes (e.g., $\nu = \sigma = 1$) ( Fig. S1(a)), the general model predicts a stronger preference for averaging compared to the simplified case, reflecting the additional contribution of signal–noise interaction terms.

### SUPPLEMENTARY FIGURES

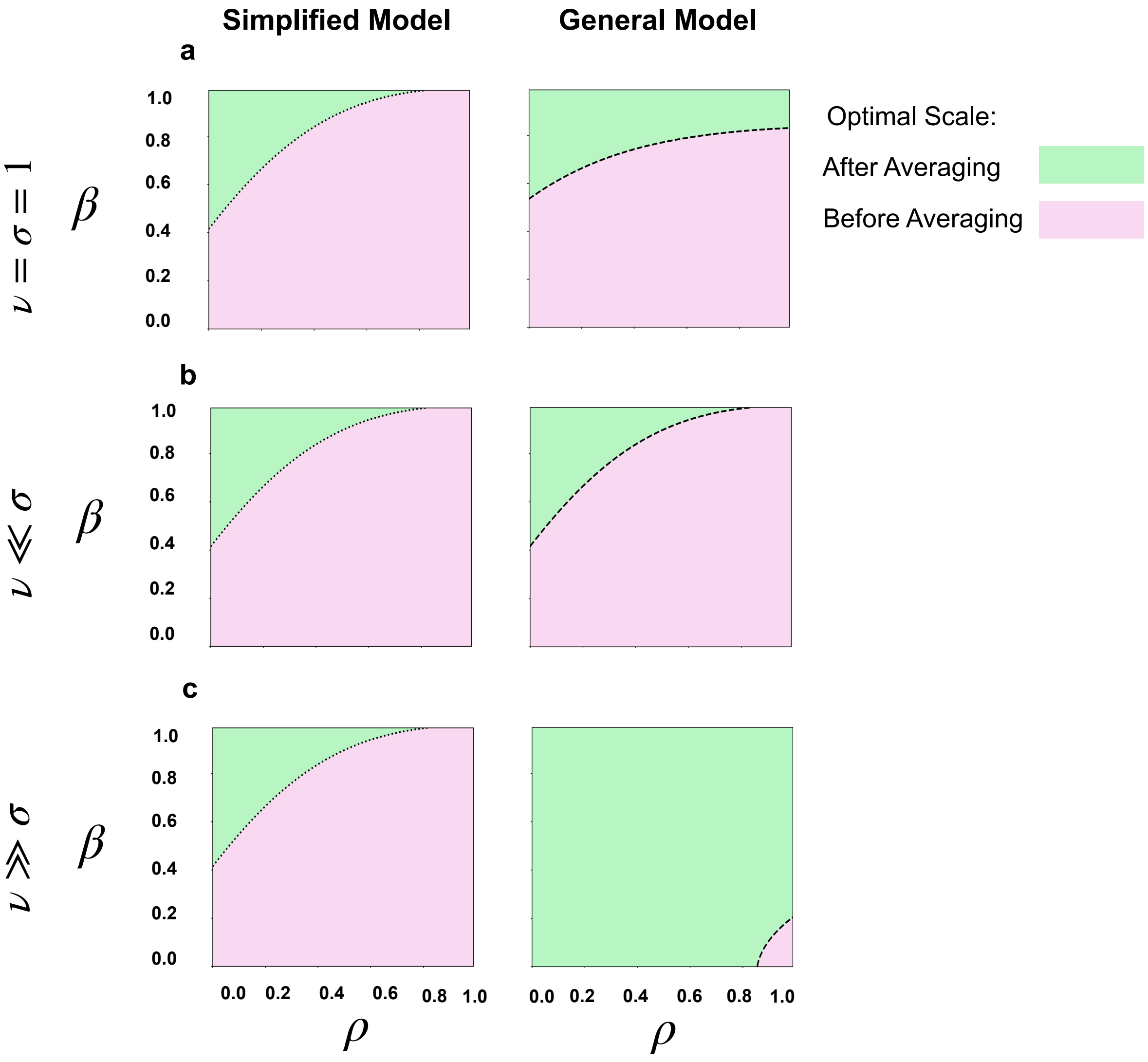


Figure S1: **The effect of signal–noise correlations on the optimal resolution of neural representations.** Each panel shows the zero-level set and corresponding signed regions of the functions $\mathcal{G}_2^{\mathrm{aft>bef}}$ in Eq. (S65) ('Simplified Model', left) and $\mathcal{H}_2^{\mathrm{aft>bef}}$ in Eq. (S66) ('General Model', right), respectively. **(a)** Comparison for the special case when $\nu = \sigma = 1$, i.e., when magnitude of signal ($\nu$, cf. Eq. (19)) equals the standard deviation of the noise ($\sigma$, cf. Eq. (5)). **(b,c)** Similar comparisons for the cases where $\nu \ll \sigma$ and $\sigma \ll \nu$, respectively. Colors represent the scale with the highest sensitivity index, as identified in the inset.

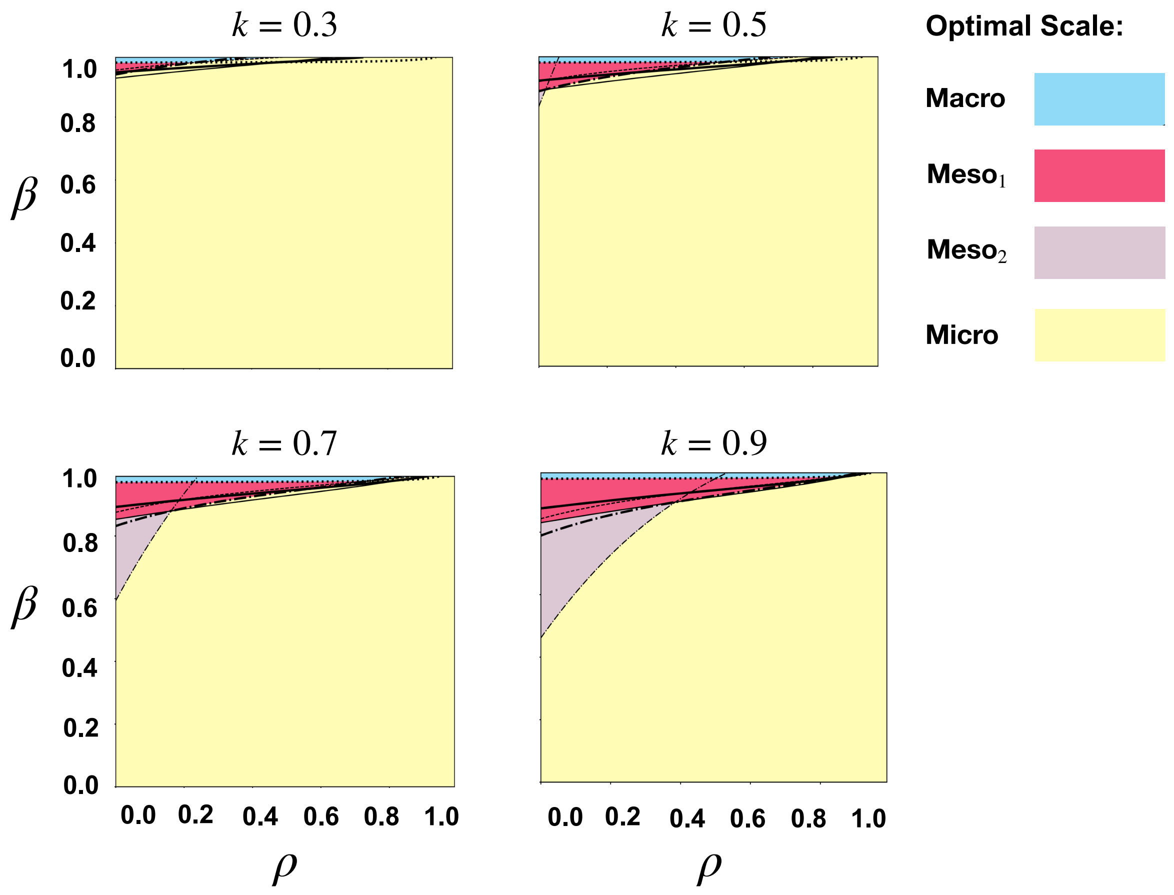


Figure S2: **Effect of the signal-autocorrelation scaling parameter $k$ on the optimal temporal resolution.** Each panel shows the global optimal-scale comparisons for increasing values of the scaling parameter $k$ and a fixed trial length of $b = 100$. Each panel overlays all six pairwise comparison boundaries. Increasing $k$ shifts the pairwise boundaries toward finer temporal scales, thereby expanding the parameter regimes in which temporal integration is beneficial. Consequently, the mesoscale-optimal region increases with increasing values of $k$. Therefore, the optimal temporal resolution continues to be determined by the balance between signal and noise autocorrelations, with arbitrary values of $k$ primarily modulating the size of the corresponding optimality regions rather than altering the overall behavior.

### SUPPLEMENTARY TABLES FOR OPTIMALITY CONDITION FUNCTIONS

Here we provide the explicit analytical forms of the optimality condition functions. These functions are obtained by substituting the closed-form expressions of the sensitivity index $d'$ into pairwise comparisons across temporal scales (microscale, mesoscale, and macroscale), followed by algebraic simplification of the resulting inequalities. Each function characterizes the relative dominance between two temporal scales. In particular, the sign of the function determines the optimal regime: a positive value indicates that the first scale yields higher discriminability, while a negative value indicates that the second scale is preferred.

Table S1 presents the optimality condition functions for the case of persistent signal correlations, where signal remains temporally aligned while noise correlations decay. Similarly, Table S2 presents the corresponding optimality conditions for the general setting in which both signal and noise correlations decay with temporal separation. In this regime, the functions incorporate geometric sums over the correlation parameters, capturing the trade-off between signal coherence and noise accumulation across scales.

| **Optimality Condition function** | **Definition** |
|---|---|
| $\mathcal{F}_b^{\mathrm{mes}_1>\mathrm{mic}}(\rho,\beta)$ | $\left[1+\left(\frac{b}{2}-1\right)\beta\right]^2-\frac{2\left[\left(\frac{b}{2}+2G\right)^2+\rho^2\left(\frac{1-\rho^{b/2}}{1-\rho}\right)^4\right]}{b+2E}$ |
| $\mathcal{F}_b^{\mathrm{mac}>\mathrm{mic}}(\rho,\beta)$ | $(1+(b-1)\beta)^2-\frac{(b+2F)^2}{b+2E}$ |
| $\mathcal{F}_b^{\mathrm{mac}>\mathrm{mes}_1}(\rho,\beta)$ | $\left[\frac{1+(b-1)\beta}{1+\left(\frac{b}{2}-1\right)\beta}\right]^2-\frac{(b+2F)^2}{2\left[\left(\frac{b}{2}+2G\right)^2+\rho^2\left(\frac{1-\rho^{b/2}}{1-\rho}\right)^4\right]}$ |
| $\mathcal{F}_b^{\mathrm{mes}_2>\mathrm{mic}}(\rho,\beta)$ | $(1+\beta)^2-\frac{2(1+\rho)^2(b+\rho^2H)}{b+2E}$ |
| $\mathcal{F}_b^{\mathrm{mes}_1>\mathrm{mes}_2}(\rho,\beta)$ | $\left[\frac{1+(\frac{b}{2}-1)\beta}{1+\beta}\right]^2-\frac{\left[\left(\frac{b}{2}+2G\right)^2+\rho^2\left(\frac{1-\rho^{b/2}}{1-\rho}\right)^4\right]}{(1+\rho)^2(b+\rho^2H)}$ |
| $\mathcal{F}_b^{\mathrm{mac}>\mathrm{mes}_2}(\rho,\beta)$ | $\frac{1+(b-1)\beta}{1+\beta}-\frac{b+2F}{(1+\rho)\sqrt{2(b+\rho^2H)}}$ |

**Table S1.** Optimality Conditions and Comparison Inequalities Across Temporal Scales for persistent signal correlations

| **Optimality Condition function** | **Definition** |
|---|---|
| $\mathcal{G}_b^{\text{mes}>\text{mic}}(\rho,\beta)$ | $\left[\frac{b(1-\beta^2)-4\beta(1-\beta^{b/2})}{b(1-\beta)^2}\right]^2 - \frac{2\left[\left(\frac{b}{2}+2G\right)^2+\rho^2\left(\frac{1-\rho^{b/2}}{1-\rho}\right)^4\right]}{b+2E}$ |
| $\mathcal{G}_b^{\text{mac}>\text{mic}}(\rho,\beta)$ | $\left[\frac{b(1-\beta^2)-2\beta(1-\beta^{b})}{b(1-\beta)^2}\right]^2 - \frac{(b+2F)^2}{(b+2E)}$ |
| $\mathcal{G}_b^{\text{mac}>\text{mes}}(\rho,\beta)$ | $\frac{b(1-\beta^2)-2\beta(1-\beta^{b})}{b(1-\beta^2)-4\beta(1-\beta^{b/2})} - \frac{b+2F}{\sqrt{2\left[\left(\frac{b}{2}+2G\right)^2+\rho^2\left(\frac{1-\rho^{b/2}}{1-\rho}\right)^4\right]}}.$ |
| $\mathcal{G}_b^{\text{mes}_2>\text{mic}}(\rho,\beta)$ | $(1+\beta)^2 - \frac{2(1+\rho)^2(b+\rho^2 H)}{b+2E}$ |
| $\mathcal{G}_b^{\text{mes}_1>\text{mes}_2}(\rho,\beta)$ | $\left[\frac{b(1-\beta^2)-4\beta(1-\beta^{b/2})}{b(1-\beta)^2(1+\beta)}\right]^2 - \frac{\left[\left(\frac{b}{2}+2G\right)^2+\rho^2\left(\frac{1-\rho^{b/2}}{1-\rho}\right)^4\right]}{(1+\rho)^2\,(b+\rho^2 H)}$ |
| $\mathcal{G}_b^{\text{mac}>\text{mes}_2}(\rho,\beta)$ | $\frac{b(1-\beta^2)-2\beta(1-\beta^{b})}{b(1-\beta)^2(1+\beta)} - \frac{b+2F}{(1+\rho)\sqrt{2b+\rho^2 H}}.$ |

**Table S2.** Optimality Conditions and Comparison Inequalities Across Temporal Scales for temporally decreasing signal correlations
